# Sphingosine kinase 1 is integral for elastin deficiency-induced arterial hypermuscularization

**DOI:** 10.1101/2024.07.01.601150

**Authors:** Junichi Saito, Jui M. Dave, Eunate Gallardo-Vara, Inamul Kabir, George Tellides, Robert K. Riemer, Zsolt Urban, Timothy Hla, Daniel M. Greif

## Abstract

Defective elastin and smooth muscle cell (SMC) accumulation characterize both arterial diseases (e.g., atherosclerosis, restenosis and supravalvular aortic stenosis [SVAS]), and physiological ductus arteriosus (DA) closure. Elastin deficiency induces SMC hyperproliferation; however, mechanisms underlying this effect are not well elucidated. Elastin (ELN) is expressed from embryonic day (E) 14 in the mouse aorta. Immunostains of *Eln(+/+)* and *Eln(-/-)* aortas indicate that SMCs of the *Eln* null aorta are first hyperproliferative at E15.5, prior to morphological differences. Bulk RNA-seq reveals that sphingosine kinase 1 (Sphk1) is the most upregulated transcript in *Eln(-/-)* aortic SMCs at E15.5. Reduced ELN increases levels of transcription factor early growth response 1 (EGR1), resulting in increased SPHK1 levels in cultured human aortic SMCs and in the mouse aorta at E15.5 and P0.5. Aortic tissue from Williams-Beuren Syndrome patients, who have elastin insufficiency and SVAS, also has upregulated SPHK1 expression. SMC-specific *Sphk1* deletion or pharmacological inhibition of SPHK1 attenuates SMC proliferation and mitigates aortic disease, leading to extended survival of *Eln(-/-)* mice. In addition, EGR1 and SPHK1 are increased in the wild-type mouse DA compared to adjacent descending aorta. Treatment with a SPHK1 inhibitor attenuates SMC proliferation and reduces SMC accumulation, leading to DA patency. In sum, SPHK1 is a key node in elastin deficiency-induced hypermuscularization, and inhibiting this kinase may be a therapeutic strategy for SVAS and select congenital heart diseases in which a patent DA maintains circulation.

**One Sentence Summary:** Sphingosine kinase 1-induced by defective elastin promotes muscularization in pathological aortic stenosis and physiological ductus arteriosus occlusion.

## INTRODUCTION

Elastin is the major component of circumferential elastic lamellae that alternate with rings of smooth muscle cells (SMCs) to form lamellar units in arteries. In humans, heterozygous loss-of-function mutations in the elastin gene *ELN* cause supravalvular aortic stenosis (SVAS) (*1*), which is characterized by aortic SMC accumulation and subsequent lumen obstruction (*2, 3*). SVAS occurs as an isolated entity or more commonly as part of Williams-Beuren Syndrome (WBS), resulting from continuous deletion of 26-28 genes including *ELN* on chromosome 7 (*4, 5*). Defective elastic lamellae and excess SMC accumulation are also observed during physiological closure of the ductus arteriosus (DA) (*6–9*). Failure of DA closure (i.e., patent DA [PDA]) leads to pulmonary congestion, systemic hypotension, and death. SMC accumulation is essential for postnatal DA closure and thus, promoting SMC proliferation may be therapeutic for PDA. In contrast, SMC accumulation is detrimental for patients with SVAS and some congenital heart diseases in which PDA is required for maintaining pulmonary or systematic circulation. Although regulating SMC accumulation is desired in these elastin-defective arteries, mechanistic links between defective elastic lamellae and SMC hyperproliferation in SVAS and DA are poorly elucidated.

Similar to SVAS patients, late-stage embryonic or early postnatal *Eln(-/-)* mice have increased vascular wall cellularity and arterial lumen obstruction, contributing to early postnatal death (*10*). *Eln(+/-)* mice have a milder aortic phenotype with a modest increase in SMC layers and thin elastic lamellae (*2, 11*). We previously demonstrated that both integrin β3 and Notch3 pathways are induced in elastin mutant mice, leading to SMC hyperproliferation (*12–14*). Although SMC hyperproliferation is the central to the pathogenesis of elastin aortopathy, it has not yet been elucidated when this excess proliferation commences.

Sphingolipids are derivatives of sphingosine, a component of the plasma membrane lipid bilayer, and play key roles in vascular biology (*15–17*); however, studies of sphingolipid signaling in elastin aortopathy or DA biology are lacking. The sphingosine kinase (SPHK) family includes SPHK1 and SPHK2 (*18*). SPHK phosphorylates sphingosine into a bioactive sphingolipid, sphingosine-1-phosphate (S1P), which binds a family of five high-affinity G-protein–coupled receptors (S1P receptors [S1PRs] 1–5) to initiate diverse intracellular events (*15–17*). Studies of pulmonary hypertension (PH) demonstrate that SPHK1, but not SPHK2, is upregulated in the lungs and pulmonary arterial SMCs from PH patients and in lungs of rodent models of hypoxia-induced PH (*19*). A recent study indicates that *Tagln-Cre, Sphk1(flox/flox)* mice have attenuated hypoxia-induced PH (*20*).

Our current study demonstrates that SMC hyperproliferation is first observed at embryonic day (E) 15.5, prior to morphological differences in the mouse aorta. Bulk RNA-sequencing (seq) indicates that Sphk1 is the most upregulated transcript in elastin mutant aortic SMCs at E15.5. Reduced ELN increases levels of SPHK1 in cultured human and mouse aortic SMCs, in elastin mutant mouse aorta at E15.5 and postnatal day (P) 0.5, in wild-type mouse DA at P0.5, and in aortic tissues from *ELN*-deficient patients. SMC-specific *Sphk1* deletion attenuates SMC proliferation, mitigating aortic disease. Similarly, pharmacological inhibition of SPHK1 attenuates SMC proliferation and hypermuscularization in elastin-defective arteries.

Treatment with the SPHK1 inhibitor extends the viability of *Eln(-/-)* mice and maintains DA patency in wild-type mice. These findings suggest that inhibiting SPHK1 is a potential therapeutic strategy for SVAS and select congenital heart diseases in which PDA maintains circulation.

## RESULTS

### Sphk1 is upregulated before morphological differences develop in elastin deficient aorta

Elastin is expressed as the monomer tropoelastin from embryonic day (E) 14 (*21*). Our initial studies are in accordance with the literature (*10*) showing that aortic medial thickness and lumen area do not differ between *Eln(+/+)* and *Eln(-/-)* mice at E13.5 and E15.5 (Fig. 1A, B). At P0.5, the *Eln(-/-)* aorta has a thick medial wall and narrow lumen (Fig. 1A, B) and an increased percent of SMCs that express the proliferation marker Ki67 (Fig. 1A-C). There is no difference in Ki67^+^ SMCs at E13.5 (Fig. 1A, C), but interestingly we observed more Ki67^+^ SMCs in the *Eln(-/-)* aorta compared to the *Eln(+/+)* aorta at E15.5, prior to a morphological phenotype (Fig. 1A-C).

**Figure 1.**
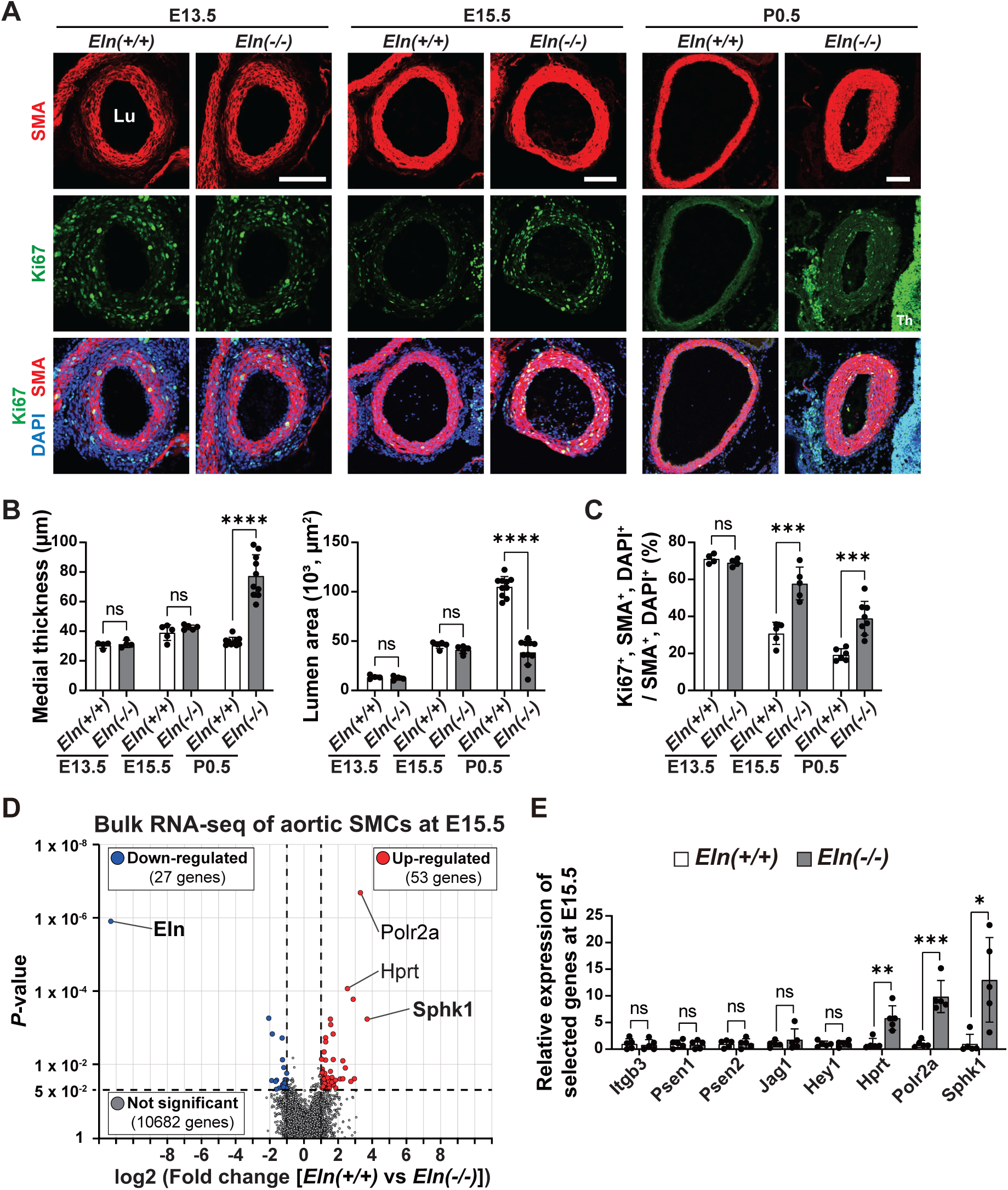
Sphk1 is upregulated before morphological differences develop in elastin deficient aorta. **(A)** Transverse sections of the ascending aorta from pups of *Eln(+/+)* or *Eln(-/-)* mice at E13.5, E15.5, and P0.5 were stained for markers of SMCs (α-smooth muscle actin [SMA]), proliferation (Ki67), and nuclei (DAPI). Lu, Lumen; Th, Thymus. Scale bars, 100 μm. **(B)** Histograms represent aortic medial wall thickness and lumen area from sections as in **A**. n=4-10 mice. **(C)** Percent of SMCs that are Ki67^+^ from sections represented in **A**. n=4-8 mice. **(D)** Volcano plot of bulk RNA-seq data of aortic SMCs isolated from *Eln(-/-)* vs *Eln(+/+)* embryos at E15.5. Genes with *P* < 0.05 and log2 fold change ≥ |1| are highlighted in red and blue color. n=5. **(E)** Expression of select genes from bulk RNA-seq data. n=5. ns, not significant, \**P* < 0.05, \*\**P* < 0.01, \*\*\**P* < 0.001 by Student’s *t* test.

Next, aortic SMCs were isolated by fluorescence-activated cell sorting (FACS) from *Acta2-GFP* (*22*) embryos carrying *Eln(+/+)* or *Eln(-/-)* at E15.5 to evaluate molecular differences (Fig. 1D). *Acta2* encodes the SMC marker protein α-smooth muscle actin (SMA). Isolated GFP^+^, CD31^-^ SMCs were subjected to bulk RNA-seq (Fig. 1D). (CD31 is an endothelial cell [EC] marker.) Bulk RNA-seq reveals significantly 53 upregulated genes and 27 down-regulated genes in *Eln* null aortic SMCs (Fig. 1D, *P* < 0.05 and log2 fold change ≥ |1|). Among the upregulated transcripts, sphingosine kinase 1 (Sphk1) is the most upregulated in *Eln(-/-)* aortic SMCs at E15.5 (Fig. 1D, E). We previously reported that integrin β3 or Notch3 (involving Jag1, Psen1/2, and Hey1) are induced in elastin mutant aortic SMCs at P0.5 (*12–14*), leading to SMC hyperproliferation. Despite excess proliferation of *Eln(-/-)* aortic SMCs at E15.5, bulk RNA-seq does not show differential expression of these genes at this early time point (Fig. 1E). Sphingolipids modulate several pathways, including SPHK1-induced activation of Notch3 in SMCs (*23, 24*), and our data support the hypothesis that Sphk1 plays a key role in the initiation of elastin aortopathy.

### Elastin insufficiency upregulates SPHK1 in mouse aorta and human aortic SMCs

To explore this hypothesis, aortas were isolated from *Eln(+/+)*, *Eln(+/-)* or *Eln(-/-)* mice at E15.5 and P0.5, and SPHK1 expression levels were analyzed in aortic lysates by quantitative real-time reverse transcription PCR (qRT-PCR) (Fig. 2A) and Western blotting (Fig. 2B).

**Figure 2.**
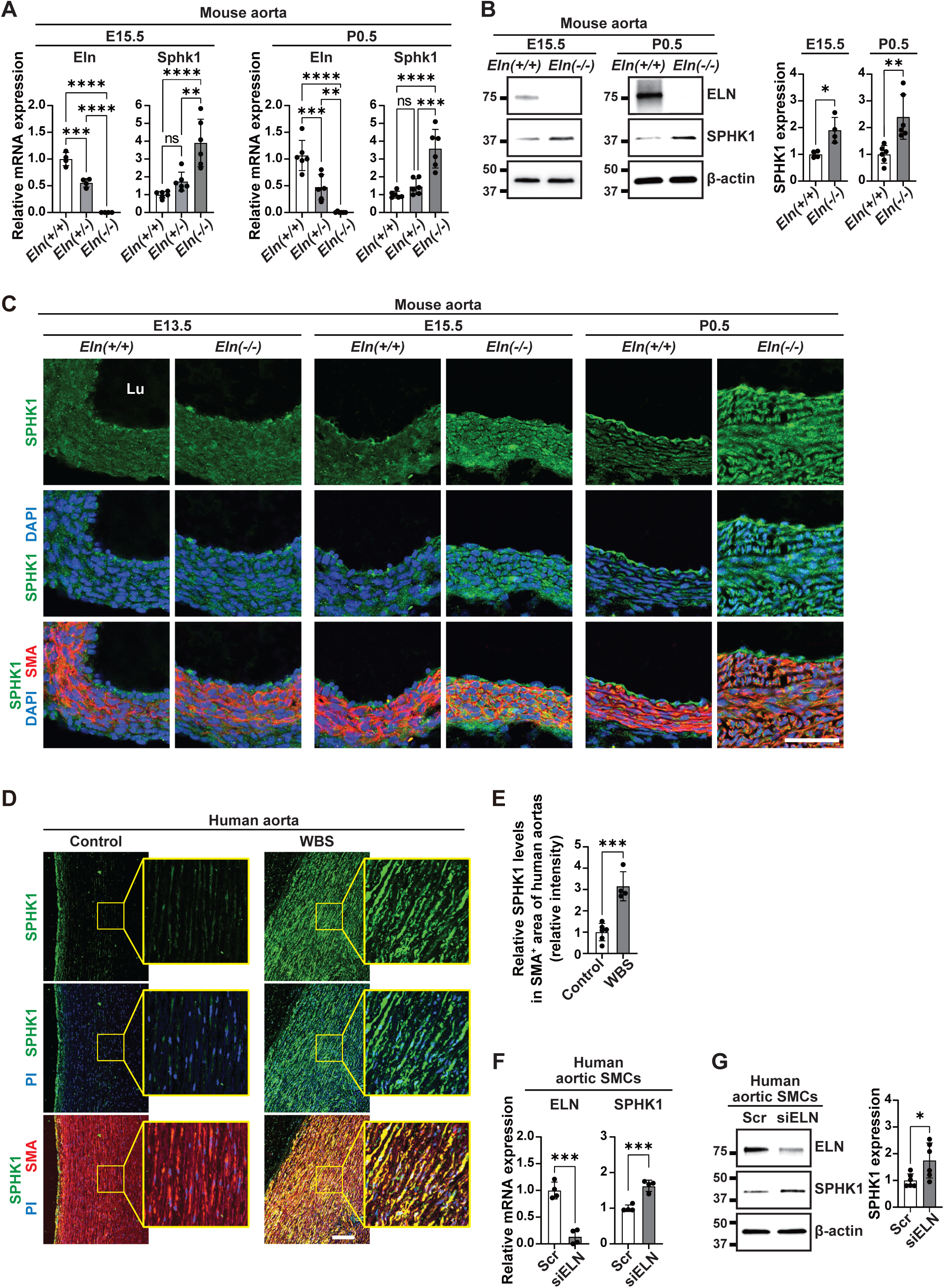
Elastin insufficiency upregulates SPHK1 in mouse and human aortic SMCs. **(A, B)** Lysates from *Eln(+/+)*, *Eln(+/-)* or *Eln(-/-)* aorta at E15.5 or P0.5 were subjected to qRT-PCR (**A**) or Western blot (**B**) for Eln or Sphk1. Densitometry of SPHK1 protein bands relative to β-actin is shown in **B**, right. n=4-6. **(C)** Transverse sections of ascending aorta from *Eln(+/+)* or *Eln(-/-)* pups at E13.5, E15.5 and P0.5 were stained for SPHK1, SMA (SMCs), and nuclei (DAPI). n=4 mice. Lu, lumen. **(D)** Human aortic sections from control male (46 years old) and Williams-Beuren Syndrome (WBS) male patient (45 years old) were stained for SPHK1, SMA, and nuclei (PI). **(E)** Relative SPHK1 levels in SMA^+^ area as in **D**. n=6 control and n=4 WBS aortas (see Table S1). **(F, G)** Lysates from human aortic SMCs treated with scrambled (Scr) or siELN were subjected to qRT-PCR (**F**) or Western blotting (**G**) for ELN and SPHK1. Densitometry of SPHK1 protein bands relative to β-actin is shown in **G**, right. n=4-6. ns, not significant, \**P* < 0.05, \*\**P* < 0.01, \*\*\**P* < 0.001, \*\*\*\**P* < 0.0001 by multifactor ANOVA with Tukey’s *post hoc* test (**A**) or Student’s *t* test (**B, E-G**). Scale bars, 50 μm (**C**), 200 μm (**D**).

Compared to *Eln(+/+)* aorta, *Eln(-/-)* aorta has higher levels of Sphk1 mRNA (∼4-fold at E15.5, and ∼3.5-fold at P0.5) and protein (∼2.0-fold at E15.5 and P0.5) (Fig. 2A, B). Transverse ascending aorta sections from *Eln(+/+)* and *Eln(-/-)* mice at E13.5, E15.5, and P0.5 were stained for SPHK1, showing upregulation at E15.5 and P0.5 with elastin deficiency (Fig. 2C). In contrast to upregulated SPHK1 levels at P0.5, SPHK2 was not changed by the reduced elastin dosage (Fig. S1). Similar to *in vivo* studies, cultured aortic SMCs isolated from *Eln(+/+)*, *Eln(+/-)*, or *Eln(-/-)* pups at P0.5 are SMA^+^ and express increased levels of Sphk1 mRNA with reduced *Eln* gene dosage (Fig. S2). Furthermore, immunohistochemical stains of aortas from patients with Williams-Beuren Syndrome (WBS) and age-matched controls (Table S1) revealed that SPHK1 is upregulated in SMCs (Fig. 2D, E). Similarly, treatment of human aortic SMCs with *ELN*-specific silencing RNA (siELN) increases SPHK1 levels (Fig. 2F, G). Taken together, these data demonstrate that elastin insufficiency upregulates SPHK1 in aortic SMCs.

### *Sphk1* deletion in SMCs attenuates elastin aortopathy

We next evaluated the effect of *Sphk1* deletion in SMCs on elastin aortopathy. *Sphk1(flox/flox)* pups that also carry no Cre or the inducible *Acta2-CreER^T2^* and either *Eln(+/+)*, *Eln(+/-)* or *Eln(-/-)* were injected daily with tamoxifen and progesterone from E13.5 to E17.5 (*12, 13*) to delete *Sphk1* in SMCs (Figs. 3A, S3). At P0.5, ascending aortas were sectioned transversely and stained for CD31, SMA, and nuclei (DAPI). In *Sphk1(flox/flox)*, *Eln(-/-)* mice, the presence of *Acta2-CreER^T2^* markedly attenuates elastin aortopathy, leading to ∼50% reduction in medial thickness and greater than ∼3-fold increase in lumen area (Fig. 3B, C).

**Figure 3.**
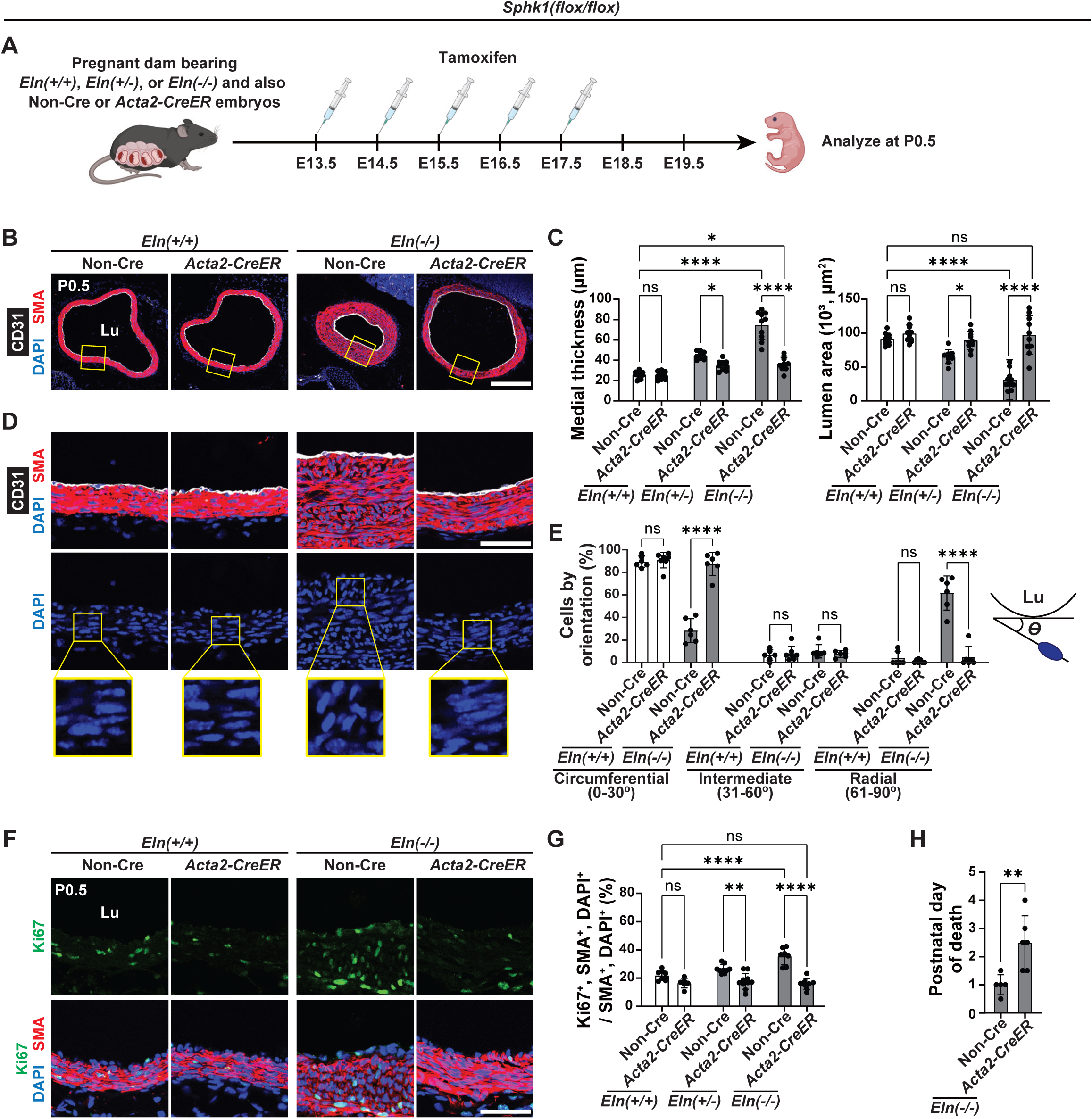
*Sphk1* deletion with *Acta2-CreER^T2^* attenuates elastin aortopathy. **(A)** Experimental strategy for (**B-H**). Pregnant dams were injected with tamoxifen daily from E13.5 to E17.5, and pups were collected at P0.5. **(B)** Transverse aortic sections from mice of indicated genotypes were stained for markers of SMCs (SMA), endothelial cells (ECs; CD31), and nuclei (DAPI). **(C)** Histograms represent aortic medial thickness and lumen area from sections as shown in **B**. n=9-10 mice. **(D)** Magnified images of **B**. **(E)** Nuclear orientation of SMCs at P0.5 with respect to tangent of the lumen boundary from images as in **D**; n=5-6 mice, and >200 SMCs were scored for each genotype. **(F)** Transverse aortic sections were stained for proliferation marker Ki67, SMA, and nuclei (DAPI). **(G)** Histograms represent percent of SMCs that are Ki67^+^ as in **F**. n=6-10 mice. **(H)** The age of death of *Eln(-/-)* pups with each genotype. n=5-6 mice. Lu, lumen. ns, not significant, \**P* < 0.05, \*\**P* < 0.01, \*\*\**P* < 0.001, \*\*\*\**P* < 0.0001 by multifactor ANOVA with Tukey’s *post hoc* test (**C, E, G**) or Student’s *t* test (**H**). Scale bars, 200 μm (**B**), 50 μm (**D, F**).

Radial misorientation of aortic SMCs in *Eln(-/-)* newborns (*13*) is attenuated by *Sphk1* deletion (Fig. 3D, E). Immunostaining for Ki67 and cleaved caspase-3 (apoptosis marker) indicates that *Sphk1* deletion in SMCs reduces elastin deficiency-induced SMC proliferation (by ∼50%; Fig. 3F, G) but does not induce apoptosis (Fig. S4). In addition, cultured elastin mutant SMCs are hyperproliferative (*25*) (Fig. S5A, B), and siSphk1 treatment reduces proliferation and wound healing (Fig. S5C-G). *Sphk1* deletion in SMCs extends the viability of *Eln(-/-)* mice (Fig. 3H), yet the extension is only ∼1.5 days despite markedly rescuing the aortic phenotype (see Fig. 3B, C). This increased survival is modest likely because *Eln(-/-)* pups have severe emphysema (*26*), and *Sphk1* deletion in SMCs does not prevent this phenotype (Fig. S6).

At P0.5, SPHK1 is expressed in ECs of the *Eln(+/+)* aorta and in both ECs and SMCs of the *Eln(-/-)* aorta (see Fig. 2C). To query the role of SPHK1 in ECs, *Sphk1(flox/flox)* mice that also carry no Cre or the inducible *Cdh5-CreER^T2^* and either *Eln(+/+)*, *Eln(+/+)* or *Eln(-/-)* were injected with tamoxifen and progesterone daily from E13.5 to E17.5 and analyzed at P0.5 (Fig. S7A-D). Immunostains reveal that *Sphk1* deletion in ECs modestly attenuates *Eln(-/-)* aortic phenotype (∼20% reduction in medial thickness and ∼70% increase in lumen area; Fig. S7E, F) and Ki67^+^ SMCs in the media (∼15% reduction; Fig. 7G, H). These findings suggest that EC-derived S1P may provide a relatively minor contribution to aortic pathology in elastin mutants.

### Pharmacological inhibition of SPHK1 attenuates aortic SMC proliferation, hypermuscularization in elastin mutants

With an eye towards clinical translation, pregnant dams bearing *Eln(+/+)*, *Eln(+/-)*, and *Eln(-/-)* embryos were injected with the SPHK1-selective competitive inhibitor PF-543 (*27, 28*) or vehicle from E13.5 until delivery, and transverse ascending aortic sections of the at P0.5 were stained for CD31 and SMA (Fig. 4A). Similar to *Sphk1* deletion in SMCs (see Fig. 3), PF-543 induces a ∼50% reduction in medial thickness and a ∼3-fold increase in lumen area of *Eln(-/-)* pups (Fig. 4B, C). SPHK1 inhibition attenuates misorientation of aortic SMCs (Fig. 4D, E), reduces SMC proliferation (Fig. 4F, G), and does not alter apoptosis (Fig. S8) in *Eln(-/-)* newborns. In addition to ascending aorta stenosis, tortuosity of the descending aorta – due to SMC hyperplasia and less axial stress – is a hallmark in *Eln(-/-)* mice (*14, 29, 30*). To visualize descending aorta morphology, yellow latex was injected through the left ventricle at P0.5 (Fig. 4H) (*14*). On *Eln(-/-)* background, SPHK1 inhibition attenuates aortic tortuosity without altering crown-rump length (Fig. 4H-J). Similar to *Sphk1* deletion in SMCs, treatment with the SPHK1 inhibitor does not improve emphysema (Fig. S9), and hence, survival of elastin null pups is only modestly extended (Fig. 4K).

**Figure 4.**
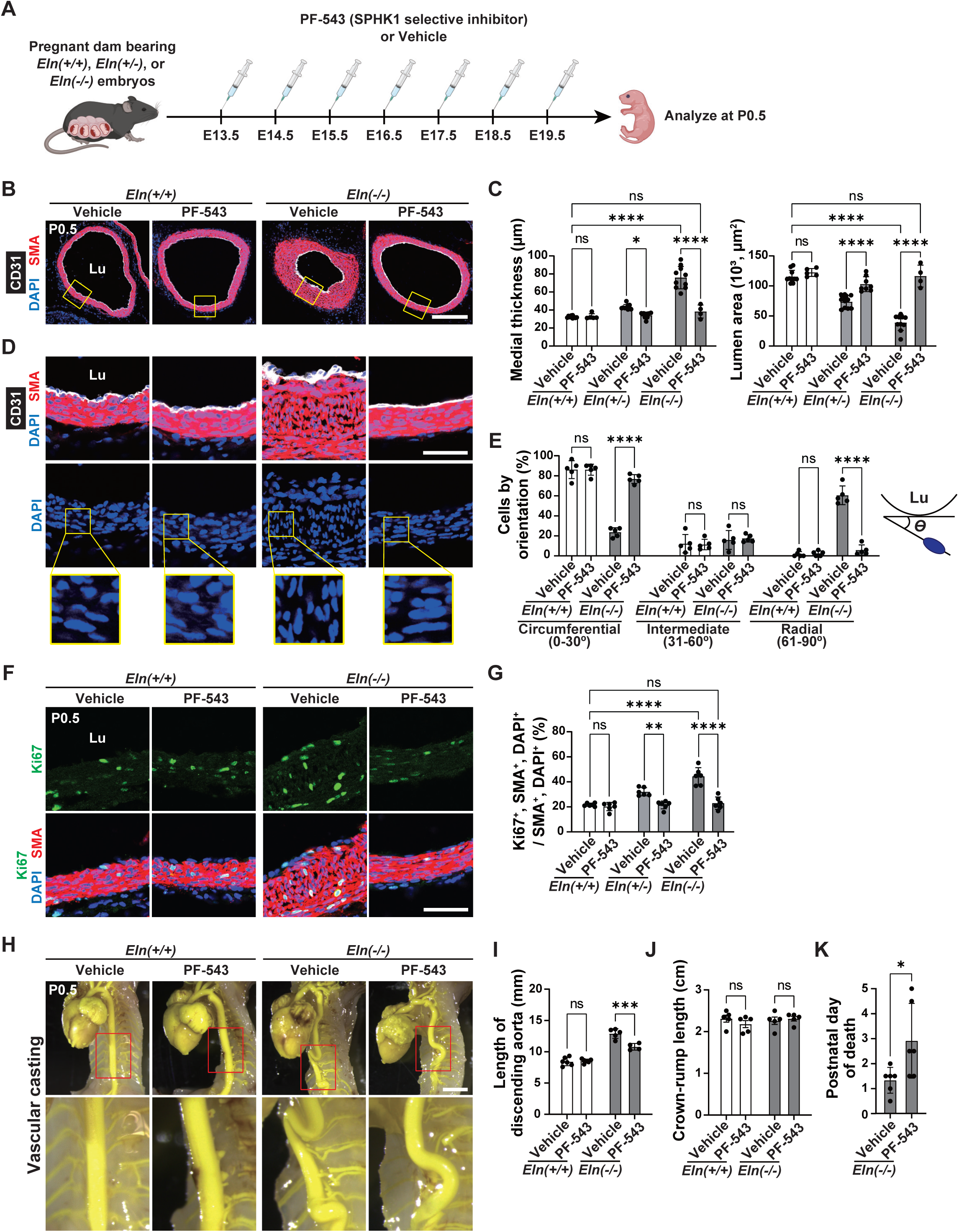
Pharmacological inhibition of SPHK1 mitigates aortopathy in elastin mutants. **(A)** Experimental strategy for (**B-K**). Pregnant dams were injected daily from E13.5-19.5 with vehicle or 20 mg/kg SPHK1 inhibitor (PF-543) (*27, 28*), and pups were collected at P0.5. **(B)** Transverse aortic sections were stained for SMA, CD31, and nuclei (DAPI). **(C)** Histograms represent aortic medial thickness and lumen area from sections as shown in **B**. n=4-10 mice. **(D)** Magnified images of **B**. **(E)** Nuclear orientation of SMCs at P0.5 with respect to tangent of the lumen boundary from images as in **D**; n=4 mice, and >200 SMCs were scored for each genotype and treatment. n=5 mice. **(F)** Transverse aortic sections were stained for Ki67, SMA, and nuclei (DAPI). **(G)** Histograms represent percent of SMCs that are Ki67^+^ as in **F**. n=6 mice. **(H)** Gross descending aorta morphology as visualized via yellow latex injections into the left ventricle of pups of indicated genotypes at P0.5 with each treatment. Lower images are magnifications. **(I)** Histograms represent length of the descending aorta from left brachiocephalic artery to celiac artery as in **H**. n=4-6 mice. **(J)** Crown-rump length of pups. n=5 mice. **(K)** The age of death of *Eln(-/-)* pups with vehicle or PF-543 treatment. n=6 mice. Lu, lumen. ns, not significant, \**P* < 0.05, \*\**P* < 0.01, \*\*\**P* < 0.001, \*\*\*\**P* < 0.0001 by multifactor ANOVA with Tukey’s *post hoc* test (**C, E, G, I, J**) or Student’s *t* test (**K**). Scale bars, 200 μm (**B**), 50 μm (**D, F**), 2 mm (**H**).

In contrast to *Eln* nulls, the lungs of *Eln(+/-)* mice develop normally (*31*) (see Fig. S6A, B) and exhibit only mild increases in both aortic medial wall thickness and SMC proliferation (*13, 14*) (Fig. S10A-C). Importantly, human SVAS/WBS, which is caused by ELN haploinsufficiency, also lacks lung parenchymal disease (*1, 2, 4*). To evaluate the postnatal therapeutic effect of SPHK1 inhibition, *Eln(+/+)* and *Eln(+/-)* pups, the latter of which develop aortopathy by P0.5, were treated with the SPHK1 inhibitor or vehicle from P2.5 to P6.5 and analyzed at P8.5 (Fig. S10D). SPHK1 inhibition significantly attenuates medial wall thickness and SMC proliferation in *Eln(+/-)* aorta (Fig. S10E-G) without inducing SMC apoptosis (Fig. S10H, I). Thus, targeting SPHK1 is a potential therapeutic strategy for SVAS/WBS patients.

### S1P receptor 1 activity is upregulated in aortic SMCs with elastin insufficiency

SPHK1 phosphorylates sphingosine to S1P, which is secreted and binds to S1P receptors [S1PRs] 1-5) to initiate cellular events including proliferation (*15*). S1PR1-3 are expressed in vascular cells (*15*), and among these receptors, S1PR1 is upregulated in *Eln(-/-)* aorta at P0.5 (Fig. S11A-C). As S1P levels in low abundant tissues (e.g., embryonic or neonatal aorta) are generally too low to be quantified, S1PR1 signaling (*S1pr1(knock-in/knock-in)*, *H2B-GFP*) mice were used to assess S1PR1 activity (*32*). Briefly, *S1pr1(knock-in/knock-in)* cells produce S1PR1 fusion protein linking the tetracycline-controlled transactivator (tTA) to its C-terminus through a tobacco etch virus (TEV) protease recognition sequence and also make a β-arrestin/TEV protease fusion protein. Upon S1P engagement to S1PR1, a β-arrestin/TEV protease is recruited to the S1PR1, resulting in the release of the tTA, which in turn induces the tTA reporter gene (i.e., *H2B-GFP*). siELN treatment of aortic SMCs isolated from *S1pr1(knock-in/knock-in)*, *H2B-GFP* mice at P0.5 upregulates Sphk1 transcript levels (Fig. S11D) and GFP expression, indicating increased S1PR1 activity (Fig. S11E, F). These data suggest that elastin insufficiency increases SPHK1 expression, leading to S1PR1 activation.

### Elastin deficiency increases Sphk1 transcripts by increasing levels of EGR1

We next addressed how elastin insufficiency upregulates Sphk1. Similar to *in vivo* data of *Eln(-/-)* aortas (see Fig. 2A), cultured *Eln(-/-)* aortic SMCs show upregulated Sphk1 transcript levels (Fig. 5A). Treatment with 5,6-Dichlorobenzimidazole 1-β-D-ribofuranoside (DRB, transcription blocker) indicates that the decay kinetics of the Sphk1 transcript is the same in *Eln(+/+)* and *Eln(-/-)* SMCs, suggesting Sphk1 transcript is not increased by reduced degradation but instead by enhanced transcription (Fig. 5B). The TRANSFAC database (https://genexplain.com/transfac/) indicates that 290 transcription factors (TFs) are reported or predicted to bind to the *SPHK1* promoter (Fig. 5C). Screening using bulk RNA-seq of cultured human aortic SMCs revealed that among 290 candidate TFs, seven TFs were upregulated by siELN (Fig. 5C). Early growth response protein 1 (Egr1) is one of the seven TFs and its upregulation with elastin deficiency is confirmed at the protein level in siELN-treated human aortic SMCs (Fig. 5D) and by qRT-PCR, Western blotting, and immunostaining of *Eln(-/-)* aorta at E15.5 and P0.5 (Fig. 5E-H). The binding of EGR1 to the *Sphk1* promoter is reported to increase SPHK1 expression in pulmonary arterial SMCs (*33*). Accordingly, silencing of Egr1 leads to a reduction of Sphk1 transcript in elastin mutant aortic SMCs (Fig. 5I). In addition, isolated murine aortic SMCs were seeded on culture plates, which were or were not pre-coated with mouse elastin (Fig. 5J), to assess the influence of elastin on Egr1 and Sphk1 levels in culture. Without coating, compared to wild-type, *Eln(-/-)* SMCs have upregulated Egr1 and Sphk1 levels (Fig. 5J). Elastin-coating significantly suppressed levels of Egr1 and Sphk1 (Fig. 5J). Thus, EGR1 is a possible transcriptional regulator of Sphk1 in elastin-deficient SMCs.

**Figure 5.**
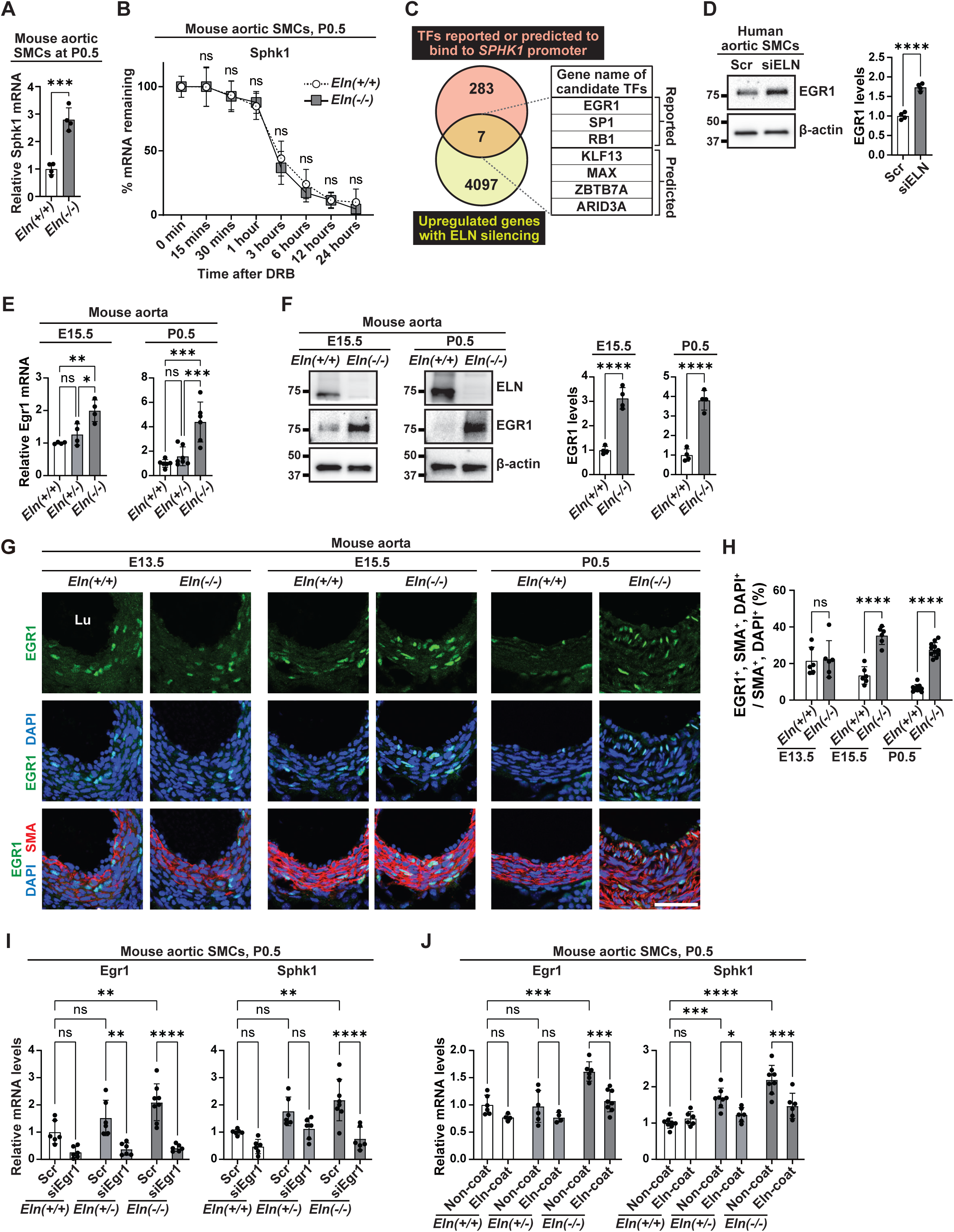
Sphk1 is upregulated by elastin deficiency-induced active transcription. **(A)** Aortic SMCs were isolated from *Eln(+/+)* or *Eln(-/-)* pups at P0.5 and subjected to qRT-PCR for Sphk1. n=4 mice. **(B)** Aortic SMCs from *Eln(+/+)* or *Eln(-/-)* pups at P0.5 were treated with transcription inhibitor DRB. mRNA expression relative to Gapdh and normalized to this ratio for time 0 is shown. n=4-10. **(C)** TRANSFAC database reports/predicts 290 transcription factors (TFs) bind to *SPHK1* promoter (-5000 bp to +100 bp). Bulk RNA-seq of human aortic SMCs with Scr or siELN (n=3 per each treatment) indicate that 7 of these 290 TFs are upregulated by ELN silencing. **(D)** Lysates of human aortic SMCs with Scr or siELN treatment were subjected to Western blotting for EGR1. Densitometry of EGR1 protein bands relative to β-actin is shown on the right. n=4. **(E, F)** Lysates from *Eln(+/+)*, *Eln(+/-)* or *Eln(-/-)* aorta at E15.5 and P0.5 were subjected to qRT-PCR (**E**) or Western blotting (**F**) for Egr1. Densitometry of EGR1 protein bands relative to β-actin is shown in **F**, right. n=4-7. **(G)** Transverse sections of ascending aorta from *Eln(+/+)* or *Eln(-/-)* pups at E13.5, E15.5 and P0.5 were stained for EGR1, SMA, and nuclei (DAPI). Lu, lumen. Scale bar, 50 μm. **(H)** Percent of SMCs that are EGR1^+^ from sections represented in **G**. n=6-11 mice. **(I, J)** Aortic SMCs isolated from *Eln(+/+)*, *Eln(+/-)*, or *Eln(-/-)* mice at P0.5 were either treated with Scr or Egr1 siRNA and cultured for 72 hours (**I**) or were cultured on elastin-coated or non-coated plates for 72 hours (**J**). Cells were then subjected to qRT-PCR for Egr1 and Sphk1. n=4-8. ns, not significant, \**P* < 0.05, \*\**P* < 0.01, \*\*\**P* < 0.001, \*\*\*\**P* < 0.0001 by multifactor ANOVA with Tukey’s *post hoc* test (**E, H-J**) or Student’s *t* test (**A, B, D, F**).

To evaluate these findings with an extracellular matrix that is potentially more relevant to physiological conditions, murine aortic SMCs from *Eln(+/+)*, *Eln(+/-)* or *Eln(-/-)* were cultured with 10% fetal bovine serum (FBS) medium for 5-10 days without changing the medium (Fig. S12A, B) (*34*). After five days of incubation, elastic fibers were partially formed by *Eln(+/+)* SMCs, whereas no elastic fibers were found in *Eln(+/-)* and *Eln(-/-)* SMC samples (Fig. S12B, left). After ten days of incubation, *Eln(+/+)* SMCs formed abundant elastic fibers, and *Eln(+/-)* SMCs deposited low levels of elastin (Fig. S12B, right). We then decellularized these samples by freeze-thaw cycles, which preserved elastic fibers (Fig. S12C, D). After decellularization, murine aortic SMCs of each *Eln* genotype were re-seeded on the resultant extracellular matrices and subjected to qRT-PCR after 48 hours of incubation (Fig. S12E). qRT-PCR revealed that *Eln(+/+)* substrates (i.e., with abundant elastic fibers) significantly attenuated Sphk1 mRNA levels in *Eln(+/-)* and *Eln(-/-)* SMCs (Fig. S12F), suggesting that extracellular matrix components (e.g., elastin) influence Sphk1 expression.

We also query the influence of mechanical properties of elastin-deficient matrix components on Sphk1 expression because the elastin-deficient aorta is stiffer than the *Eln(+/+)* aorta (*11*). It is reported that in metastatic cancer cells, stiffer substrates increase Sphk1 expression and S1P production (*35*). Our data with PDMS substrates (*36*) indicates that in *Eln(-/-)* aortic SMCs, stiffer substrates increase SPHK1 levels compared to softer substrates (Fig. S13), suggesting the involvement of substrate stiffness in Sphk1 regulation.

### SPHK1 and EGR1 are upregulated in mouse ductus arteriosus (DA)

Similar to elastin aortopathy, physiologically developed DA is characterized by defects in elastic lamellae and excess SMC accumulation (*6–9*). These changes in the vascular wall are implicated in contributing to postnatal DA closure; however, in the DA, data connecting defective elastic lamellae and SMC accumulation are lacking. To evaluate SPHK1 signaling in the DA, transverse sections of DA and the adjacent descending aorta from wild-type pups at P0.5 were stained for elastin, SPHK1, and EGR1 (Fig. 6). Consistent with previous literature (*8*), elastin staining showed fewer elastic fibers in the DA compared to the adjacent aorta (Fig. 6A, B). Inversely, SPHK1 and EGR1 are upregulated in the DA (Fig. 6C-G). Furthermore, in isolated tissues from wild-type pups at P0.5, qRT-PCR and Western blotting confirm upregulated SPHK1 and EGR1 in the DA compared to the descending aorta (Fig. 6H, I).

**Figure 6.**
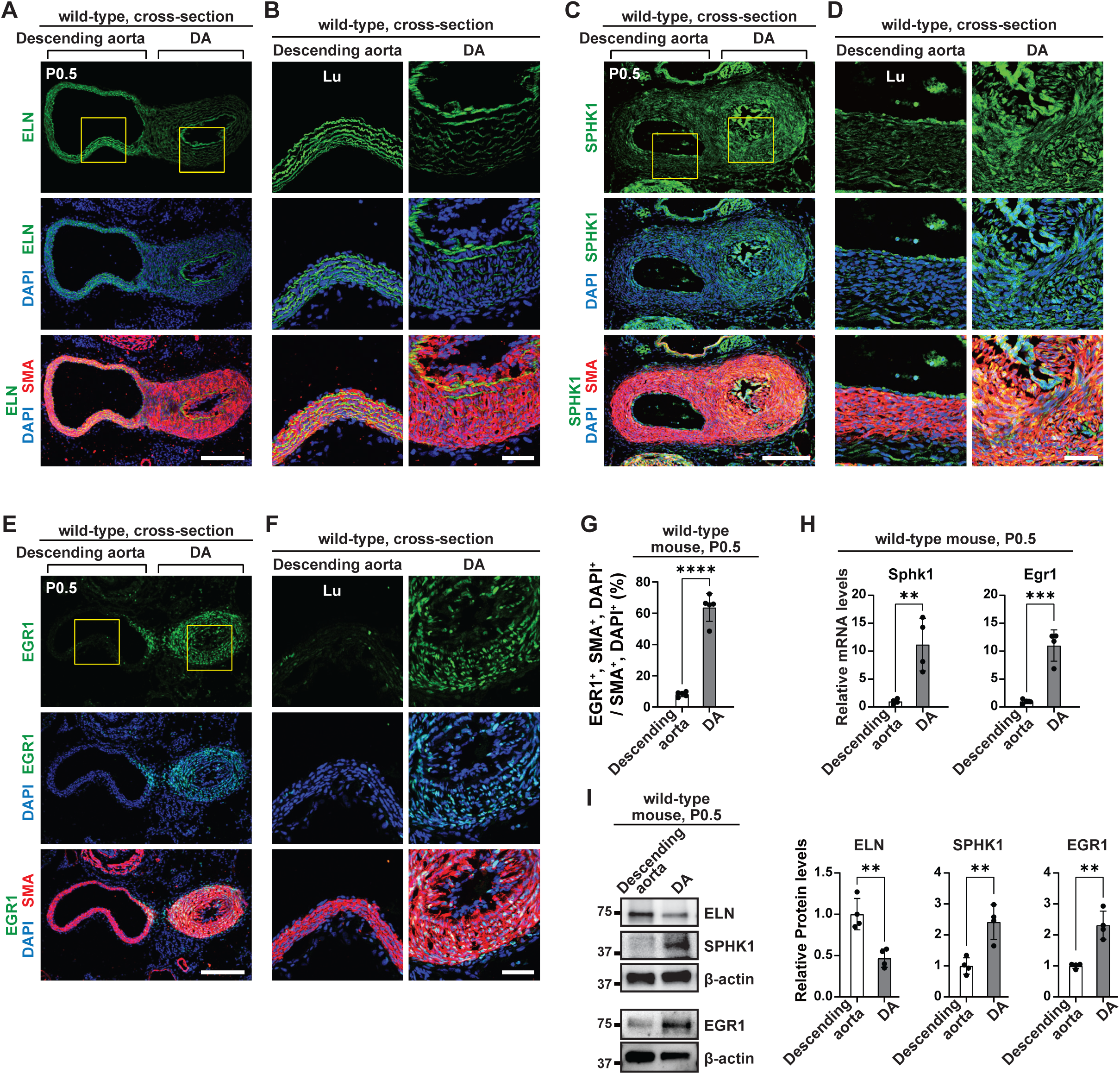
SPHK1 and EGR1 are upregulated in murine ductus arteriosus (DA). **(A-F)** DA and adjacent descending aorta cross-sections of wild-type mice at P0.5 were stained for SMA, nuclei (DAPI), and either ELN in **A, B**, SPHK1 in **C, D** or EGR1 in **E, F**. Images in **B, D, F** are magnified from **A, C, E**, respectively. Lu, lumen. **(G)** Percent of SMCs that are EGR1^+^ from sections represented in **F**. n=5 mice. **(H, I)** Lysates from wild-type DA or aorta were subjected to qRT-PCR (**H**) or Western blotting (**I**) for SPHK1, EGR1 and/or ELN as indicated. Densitometry of protein bands relative to β-actin is shown in **I**, right. n=4. \*\**P* < 0.01, \*\*\**P* < 0.001, \*\*\*\**P* < 0.0001 by Student’s *t* test. Scale bars, 200 μm (**A, C, E**), 50 μm (**B, D, F**).

### SPHK1 inhibition attenuates SMC accumulation, leading to postnatal PDA

To evaluate the effect of SPHK1 inhibition in DA biology, pregnant dams bearing wild-type embryos were injected daily with PF-543 or vehicle from E13.5 until delivery, and pups were analyzed at P0.5 (Fig. 7A). Long axis sections of the DA at P0.5 were stained for SMA, CD31 and nuclei (DAPI) (Fig. 7B, C). Newborns exposed to vehicle have prominent SMC accumulation (i.e., intimal thickening [IT]), which occludes the DA lumen; however, PF-543 treatment reduces Ki67^+^ SMCs in the DA and attenuates SMC accumulation, leading to PDA at P0.5 (Fig. 7B-F). Because of variability of the time of parturition, we further evaluated kinetics of postnatal DA closure after cesarean section at E18.5 (to standardize the time of delivery) following injection of pregnant dams with PF-543 or vehicle from E13.5-17.5 (Fig. S14A).

**Figure 7.**
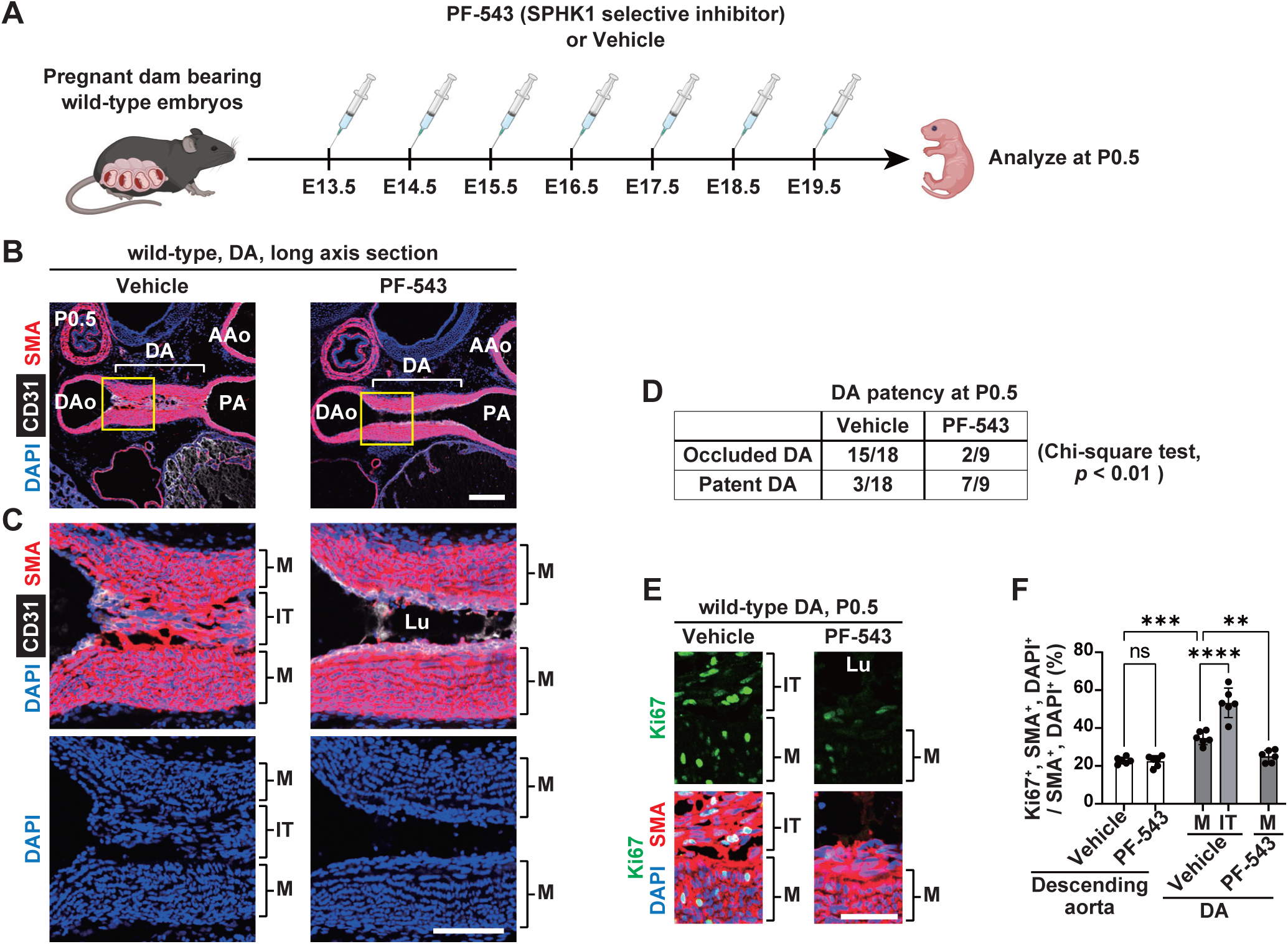
SPHK1 inhibition attenuates hypermuscularization, leading to DA patency. **(A)** Experimental strategy for (**B-F**). Pregnant dams bearing wild-type embryos were injected daily E13.5-19.5 with SPHK1 inhibitor (PF-543, 20 mg/kg) or vehicle, and pups were collected at P0.5. **(B, C)** DA sections were stained for SMA, CD31, and nuclei (DAPI). DAo, descending aorta; PA, pulmonary artery; AAo, ascending aorta. **(C)** Magnified images of **B**. M, media; IT, intimal thickening; Lu, lumen. **(D)** Numbers of occluded or patent DA at P0.5. n=9-18 mice. **(E)** Long axis DA sections were stained for Ki67, SMA and nuclei (DAPI). **(F)** Percent of SMCs that are Ki67^+^ as in **E**. n=6 mice. ns, not significant, \*\**P* < 0.01, \*\*\**P* < 0.001, \*\*\*\**P* < 0.0001 by multifactor ANOVA with Tukey’s *post hoc* test. Scale bars, 200 μm (**B**), 100 μm (**C**), 50 μm (**E**).

Newborns with either treatment showed DA constriction after birth (Fig. S14B, C). Yet compared to vehicle-treated DA, SPHK1 inhibition results in less muscularization and increased lumen area (Fig. S14C).

Available drugs for DA target vascular tone via prostaglandin E2 (PGE2) signaling and no medical treatment specifically targets SMC accumulation. To evaluate whether SPHK1 signaling is independent of PGE2 signaling, DA and aortic SMCs from *Eln(+/+)* pups at P0.5 were treated with either vehicle (0.1% DMSO), PGE2 (10^-6^ µM), or indomethacin (an inhibitor of prostaglandin synthesis; 10^-5^ µM) (*37*) and subjected to qRT-PCR (Fig. S15A). Consistent with a prior study (*38*), glycoprotein fibulin 1 (Fbln1), which regulates DA SMC migration, was increased by PGE2 and decreased by indomethacin (Fig. S15A). In contrast, Egr1 and Sphk1 transcripts were not changed by these treatments (Fig. S15A). In addition, silencing of Sphk1 does not alter Fbln1 and cyclooxygenase 2 (Cox2) transcript levels (Fig. S15B). These data suggest that elastin deficiency-induced Sphk1 signaling is independent of PGE2.

In summary, elastin insufficiency in SMCs leads to upregulation of SPHK1 through EGR1-induced transcription (Fig. 8). Enhanced SPHK1 induces S1PR1 activation and SMC hyperproliferation. Similar to elastin aortopathy, SPHK1 and EGR1 are increased during physiological closure of the DA. SPHK1 deletion and/or inhibition attenuates SMC accumulation in pathological elastin aortopathy and in physiological DA closure. Thus, inhibiting SPHK1 is an attractive therapeutic strategy for SVAS and select congenital heart diseases in which PDA is requisite to maintain circulation.

**Figure 8.**
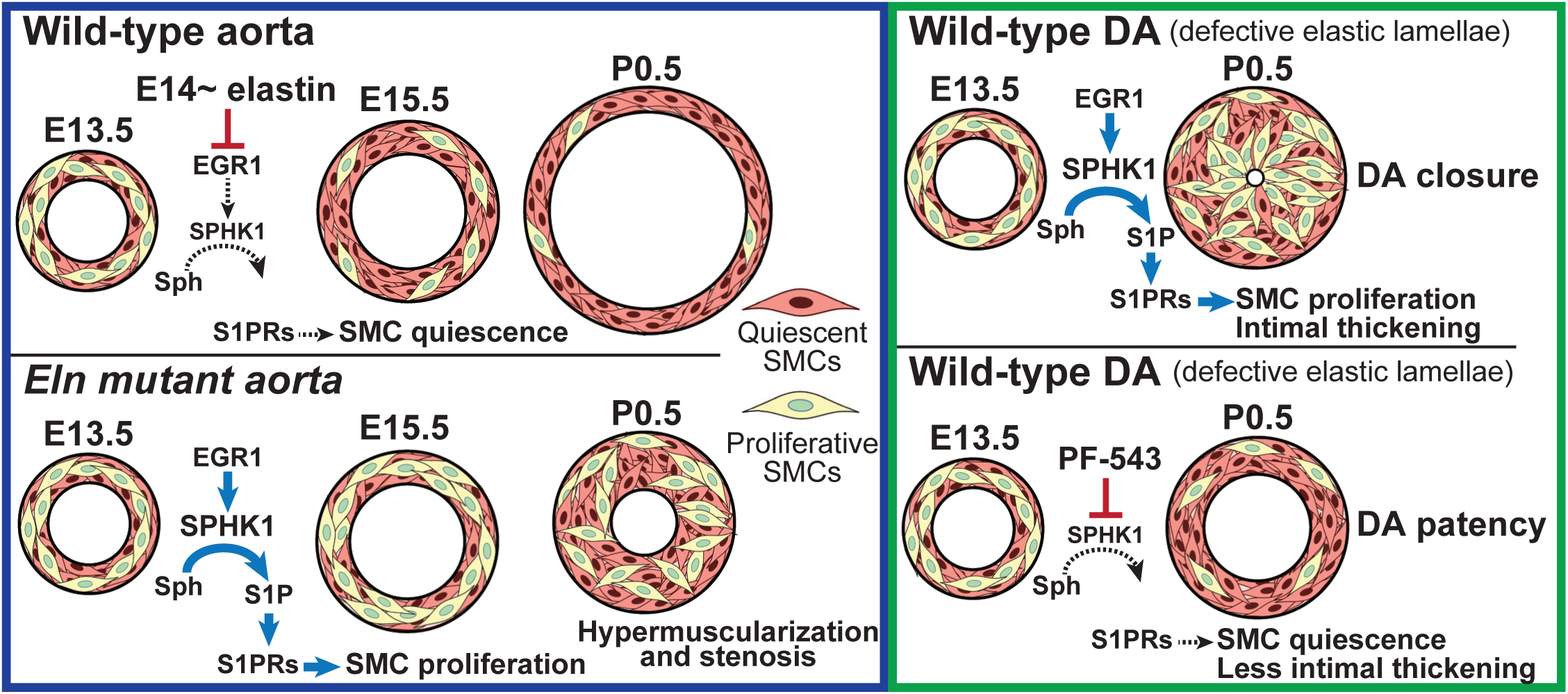
Schematic representation of working model. Defective elastin upregulates EGR1 and thereby, SPHK1. Upregulated SPHK1 phosphorylates sphingosine (Sph) into sphingosine-1-phosphate (S1P), which activates S1P receptors (S1PRs), resulting in SMC hyperproliferation. PF-543, SPHK1-selective inhibitor.

## DISCUSSION

The current study identifies SPHK1 as an early responsive molecule to elastin deficiency by analyzing SMCs at E15.5 when hyperproliferation is first observed in the *Eln(-/-)* aorta (Fig. 1). SPHK1 is generally implicated in promoting cell proliferation (*17*), whereas SPHK2 suppresses cell growth and induces apoptosis (*39, 40*). In mice, upregulated SPHK1 in the lung and heart is associated with hypoxia-induced pulmonary hypertension (*19*) and angiotensin II-induced cardiac hypertrophy (*41*), respectively. These effects are thought to result from SPHK1-derived S1P. Indeed, S1P promotes proliferation of various cell types including aortic SMCs (*42*). Both *Sphk1(-/-)* and *Sphk2(-/-)* mice are viable without significant phenotypes, but double mutants (i.e., *Sphk1(-/-)*, *Sphk2(-/-)*) exhibit dilated blood vessels and severe hemorrhage and do not survive beyond E13.5 (*18, 43*). S1P is irreversibly degraded by S1P lyase (Sgpl1), and *Sgpl1(-/-)* mice exhibit vascular developmental defects, dying within 6 weeks of birth (*44*). The proper expression/activity of SPHK1-S1P signaling is indispensable for vascular development.

The cellular response to S1P depends on expressed S1P receptor subtypes. S1PR1, S1PR2 and S1PR3 are expressed in vascular SMCs, whereas S1PR1 and S1PR3 are predominant S1P receptors in ECs (*45, 46*). Prior studies provide evidence that S1PR1 promotes, whereas S1PR2 attenuates, SMC proliferation (*42, 47–49*). Compared to adult rat vascular SMCs, rat pup intimal SMCs express higher S1pr1 transcript levels, showing a greater proliferative response to S1P (*47*). S1P-induced SMC proliferation in culture is attenuated by inhibition of S1PR1/S1PR3 and potentiated by S1PR2 inhibition (*42*). Ligation-induced SMC proliferation and neointimal hyperplasia in mouse carotid arteries are enhanced by overexpression of *S1pr1* under the *Acta2* promoter or global knockout of *S1pr2* (*48, 49*). Interestingly, S1PR1 is detected only in ECs of uninjured carotid arteries, while S1PR1 is detected both in the neointima and ECs in the injured arteries (*48*). A similar pattern was found in our data with S1PR1 mainly expressed in ECs of *Eln(+/+)* aorta, while it is upregulated in SMC layers of *Eln(-/-)* aorta (Fig. S11C). These findings indicate that aortic SMCs with elastin insufficiency have enhanced S1PR1 expression and activity (Fig. S11), resulting in SMC hyperproliferation (Figs. 1, 3, 4).

Pre-natal treatment with SPHK1 inhibitor PF-543 prevents elastin aortopathy (Fig. 4). Furthermore, postnatal PF-543 attenuates pre-established SMC proliferation and muscularization in *Eln(+/-)* aorta (Fig. S10). Other studies demonstrate the efficacy of PF-543 in rodent models of pulmonary hypertension, bronchopulmonary dysplasia, and lung fibrosis (*27, 28, 50*). However, PF-543 is not yet approved by the Food and Drug Administration, and its use is limited in part by low stability (*51*). In contrast, S1PR1 antagonists fingolimod and siponimod are approved for use as immunomodulators, primarily for multiple sclerosis (*15*). Our results provide evidence of upregulated SPHK1-S1P-S1PR1 signaling in elastin aortopathy, suggesting the therapeutic potential of inhibiting this pathway for combating SMC accumulation in SVAS.

In addition to SVAS, physiological DA closure is promoted by enhanced SPHK1 signaling (Figs. 6, 7, S14). Closure of the DA is a product of vascular constriction and SMC accumulation (*52*). Current drugs for DA target vascular tone through PGE signaling (*52*). Upon PGE binding to PGE receptor 4, which is highly expressed in DA SMCs, cyclic AMP inhibits myosin light chain kinase, resulting in DA relaxation (*53*). Inversely, treatment with cyclooxygenase inhibitors (e.g., indomethacin) prevents prostaglandin synthesis, leading to DA constriction (*53*). Studies with rodent models indicate that PGE signaling also induces SMC accumulation through hyaluronan production (*37*), enhanced focal adhesion (*54*), and increased FBLN1 expression (*38*). Thus, there is a therapeutic dilemma: PGE induces DA dilation but promotes SMC accumulation, and cyclooxygenase inhibition induces vasoconstriction but prevents SMC accumulation. The current study suggests that SPHK1 signaling is independent of PGE signaling (Fig. S15). Notably, most patients with persistent PDA need treatment for closing the DA, whereas patients with select congenital heart diseases require treatment for maintaining the PDA to sustain systemic or pulmonary circulation. Considering the independence of SPHK1 and PGE signaling in the DA, strategies of both promoting and inhibiting SPHK1 are promising for controlling PDA in combination with drugs targeting PGE signaling.

Quantity and activity of SPHK1 is regulated by transcription, translation, post-translational modifications, and degradation (reviewed in (*55*)). Our data indicate that elastin insufficiency upregulates both Egr1 transcript levels and EGR1-mediated Sphk1 transcription (Fig. 5). Thus, a further question arises: how does elastin insufficiency upregulate Egr1 transcripts? Although beyond the experimental scope of the current paper, we speculate a number of possibilities underlying Egr1 upregulation. The first possibility stems from prior studies indicating that Egr1 is mechanosensitive (*56, 57*). As *Eln(-/-)* aorta is stiffer than *Eln(+/+)* aorta (*11*), the stiff elastin-deficient matrix may induce Egr1 expression. Indeed, our data indicate that the Sphk1 transcript is upregulated by a stiff substrate (Fig. S13). Disrupted elastic fibers are also observed in mice null for the gene encoding lysyl oxidase (*Lox*) (*58*) or fibulin4 (*Fbln4*) (*59*). Interestingly, Egr1 levels are upregulated in *Lox(-/-)* aorta (*29*), and *Egr1* deletion ameliorates the formation of thoracic aortic aneurysm in SMC-specific *Fbln4(-/-)* mice (*60*). Another possible contributor is a positive feedback loop of EGR1-SPHK1-S1P signaling (*33, 48, 61*). A prior study reported that Egr1 mRNA is increased by S1P stimulation and/or S1PR1 overexpression in cultured mouse aortic SMCs (*48*). The upregulated EGR1 in turn increases SPHK1 expression and thus, leads to increased S1P production (*33, 48, 61*) (Fig. 8).

Taken together, our results highlight SPHK1 signaling as an integral connection between defective elastin and SMC hyperproliferation (Fig. 8). The consistency of this finding across different arteries in distinct contexts emphasizes the importance of SPHK1 signaling in vascular development, homeostasis, and disease. In addition to pathological SVAS and physiological DA, diverse arterial diseases including atherosclerosis, restenosis, and pulmonary hypertension, are characterized by defective elastic lamellae and hypermuscularization (*25, 62*), suggesting a broader relevance of the current work. Further studies of SPHK1 signaling are warranted and promise to advance novel therapeutic strategies for defective elastin-induced arterial muscularization.

## MATERIALS AND METHODS

### Study design

The overall objectives of this study were to identify an early responsive gene in elastin deficiency and determine the gene product’s function in elastin aortopathy and physiological DA closure. To this end, we studied elastin mutant aorta and wild-type DA from mice as well as aorta samples from humans with elastin insufficiency. For mouse studies, a power analysis with 80% power and α = 0.05 indicated that a sample size of a minimum of four mice per group was required for expression analysis and gene deletion experiments as in our prior studies (*12, 14*).

For studies with human samples, a power analysis based on our prior study (*12*) determined that samples from at least four distinct patients per group would provide 95% power to detect a significant difference with α = 0.05. The number of repeats for each experiment is included in figure legends. Endpoints were prospectively selected on the basis of our previous studies on elastin aortopathy (*12–14*). All data are included (no outliers were excluded).

### Mice

C57BL/6 wild-type mice were from The Jackson Laboratory. Mouse strains used were *Eln(+/-)* (*10*), *Acta2-GFP* (*22*), *Acta2-CreER^T2^* (*63*), *Cdh5-CreER^T2^* (*64*)*, Sphk1(flox/flox)* (*65*), *S1pr1(knock-in/knock-in)* (*32*), and *H2B-GFP* (*32*). Mice were bred, and embryos or pups were harvested at different ages with embryonic day (E) 0.5 considered noon of the day that the vaginal plug was detected.

For *Sphk1* deletion in *Sphk1(flox/flox)* mice, 1 mg tamoxifen (Sigma-Aldrich, T5648) and concomitant 0.25 mg progesterone (Sigma-Aldrich, P3972; to prevent dystocia) were administered intraperitoneally to pregnant dams daily from E13.5 to E17.5, and pups were analyzed at P0.5. For pharmacological inhibition of SPHK1, pregnant dams were intraperitoneally injected with PF-543 (20 mg/kg body weight, Cayman Chemical, 17034-25) or vehicle (20% 2-hydroxypropyl β-cyclodextrin, Sigma-Aldrich, H107) daily from E13.5 until delivery, and pups were analyzed at P0.5. In select experiments, *Eln(+/+)* or *Eln(+/-)* pups were injected each day with PF-543 (20 mg/kg body weight) from P2.5 to P6.5 and analyzed at P8.5. All mouse experiments were approved by the Institutional Animal Care and Use Committee at Yale University and in accordance with the NIH *Guide for the Care and Use of Laboratory Animals* (National Academies Press, 2011).

### Immunohistochemistry

Murine embryos or pups were euthanized and fixed in 4% paraformaldehyde for 2 hours. These samples were washed with PBS for 2 hours and incubated with 30% sucrose in PBS for 24-48 hours at 4°C until the sample sunk. Samples were embedded in OCT compound (Tissue-Tek), frozen on dry ice, and stored at –80°C. Transverse serial cryosections of ascending aortas were cut 8 μm thick starting immediately caudal to the aortic arch and ending at the aortic valve.

As sections near the aortic arch show most robust aortic phenotypes in elastin mutants (*14*), these sections were utilized for morphological analysis. Cryosections were washed with 0.5% Tween 20 in PBS (PBS-T) and incubated with blocking solution (5% goat serum, 0.1% Triton X-100 in PBS) for 1 hour and then with primary antibodies diluted in blocking solution overnight at 4°C. On the next day, sections were washed with PBS-T and incubated with secondary antibodies diluted in blocking solution for 1 hour. Primary antibodies used were rat anti-CD31 (BD Pharmigen, 553370; 1:250), rabbit anti-SPHK1 (Proteintech, 10670-1-AP; 1:200), rabbit anti-Ki67 (Invitrogen, MA5-14520; 1:250), rabbit anti-cleaved caspase-3 (Cell signaling, 9664; 1:250), rabbit anti-EGR1 (Cell signaling, 4154; 1:1000), rabbit anti-S1PR1 (Proteintech, 55133-1-AP, 1:200), chicken anti-GFP (Abcam, ab13970; 1:100), rabbit anti-fibronectin (F3648, ThermoFisher; 1:100), and directly conjugated Cy3 mouse anti-SMA (Sigma-Aldrich, C6198; 1:500). Secondary antibodies used were conjugated Alexa Fluor 488 or 647 against rat or rabbit (ThermoFisher, A11008, A11006, A21245, A21247; 1:500) and conjugated Alexa Fluor 488 against chicken (Abcam, ab150169; 1:500). DAPI (Sigma-Aldrich, D9542; 1:300) was used for nuclear staining. CF633 hydrazide was used for elastin staining (Sigma-Aldrich, SCJ4600037; 1:20000) (*12, 66*).

Aortic samples from de-identified patients with WBS and controls (see Table S1) were fixed in formalin, paraffin embedded, and sectioned. Paraffin was removed with Histo-Clear (National Diagnostics), and after ethanol washes, sections were rehydrated into water.

Rehydrated sections were incubated in boiling antigen retrieval buffer (Dako, S1699) for 20 minutes. Sections were allowed to cool at room temperature for 1 hour and then rinsed twice in PBS-T and blocked for 1 hour in 5% goat serum, 0.5% Triton X-100 in PBS. Slides were incubated overnight with anti-SPHK1 antibody (Proteintech, 10670-1-AP; 1:100) at 4 °C. The next day, sections were washed with PBS-T and incubated for 1 hour with secondary Alexa Fluor 647 conjugated antibody against rabbit (ThermoFisher, A21245; 1:500) and FITC conjugated anti-SMA antibody (Sigma-Aldrich, F3777, 1:500). Propidium iodide (Sigma-Aldrich, P4170; 1:100) was used for nuclear staining.

### Quantification of parameters of aortic morphology and staining intensity

Media and lumen parameters were quantified on transverse sections of mouse aorta with ImageJ software (NIH). Medial wall thickness was calculated by measuring the distance between the inner aspect of the inner and the outer aspect of the outer SMA^+^ medial layers (*12, 14*). Lumen area was calculated by measuring the area interior to CD31 staining. Fluorescence intensity of immunostaining for SPHK1 on formalin-fixed, paraffin-embedded human aortic sections (10 μm thick) was measured from four WBS patients and six controls (4 fields per sample). On the ImageJ software, each image was split into individual color channels, such as green for SPHK1 in Figure 2D. SMA^+^ area (200 x 200 µm^2^ in size) was then randomly selected (4 areas per image) and measured for green color intensity (i.e., SPHK1 expression). Intensity of SPHK1 staining from the aortas of WBS patients was normalized to that of age-matched controls.

### Cell isolation

Aortic SMCs were isolated from E15.5 embryos carrying *Acta2-GFP* by enzymatic digestion and FACS. As described previously (*37*), dissected aortas were incubated with collagenase-dispase enzyme mixture (1.5 mg/ml collagenase-dispase [Sigma-Aldrich, 10269638001], 0.5 mg/ml elastase [Sigma-Aldrich, E0127], 1 mg/ml trypsin inhibitor [Sigma-Aldrich, T6522], and 2 mg/ml bovine serum albumin [Sigma-Aldrich, A9418] in Hanks’ balanced salt solution [Sigma-Aldrich, H9269]) on a shaker at 37 C° for 20 minutes, and then with collagenase II enzyme mixture (1 mg/ml collagenase II [Worthington, LS004176], 0.3 mg/ml trypsin inhibitor, and 2 mg/ml bovine serum albumin in Hanks’ balanced salt solution) at 37 C° for 12 minutes. The enzymatic reaction was stopped with a 20% FBS M199 medium.

After centrifuging the cell suspensions and removing the supernatant, the pellet was resuspended in stain buffer (BD Pharmingen, 554656) and incubated with Alexa Fluor 647 conjugated CD31 antibodies (Invitrogen, A14716) for 30 minutes on ice to select CD31^-^ cells by FACS. After centrifugation and resuspension in stain buffer containing DAPI, cell suspensions were subjected to FACS. Isolated GFP^+^, CD31^-^, DAPI^-^ SMCs were incubated in RNA lysis buffer (Qiagen, 74004) and subjected to RNA extraction and bulk RNA-seq as described below.

For cell culture, SMCs were isolated from harvested aorta or DA of pups at P0.5. Adventitial layers were carefully removed with fine forceps. Aortas were then cut open, and the lumen side was repeatedly scraped with fine forceps to remove ECs. The remaining medial layers were digested using the collagenase-dispase and collagenase II enzyme mixture as described above. Digested SMCs were cultured on plates pre-coated with poly-L-lysine solution (Sigma-Aldrich, P4707). The isolated cells were confirmed to have morphological (hill-and-valley appearance) and gene product expression (SMA, checked by qRT-PCR, Western blot, and immunocytochemistry) characteristics of SMCs.

### Cell culture

Human aortic SMCs were purchased from Lonza (CC-2571), and murine SMCs were isolated as described above. SMCs were cultured up to passage 6 in M199 medium supplemented with 10% FBS, epidermal growth factor (3 μg/ml), and fibroblast growth factor (2 μg/ml) (PeproTech). In select experiments, mouse elastin (Sigma-Aldrich, E6402, 50 μg/ml) was pre-coated on the surface of cell culture plates.

### siRNA-mediated knockdown

As described previously (*12*), human or murine aortic SMCs were transfected with Lipofectamine 2000 (Life Technologies) containing Scr RNA or siRNA targeting *ELN*, *SPHK1* or *EGR1* (Horizon Discovery Biosciences Limited, 50 nM) for 6 hours. Cells were then washed in M199 medium and cultured for 72 hours prior to collection for qRT-PCR, Western blotting, and/or bulk RNA-seq.

### Drug treatment in culture

Murine SMCs isolated from the descending aorta and DA at P0.5 as described above were seeded on cell culture plates. SMCs were starved with 0.5% FBS M199 medium for 24 hours and then incubated with either vehicle (0.1% DMSO), PGE2 (Sigma-Aldrich, P0409, 10^-6^ µM), or indomethacin (Sigma-Aldrich, I7378, 10^-5^ µM) for 48 hours before collection for qRT-PCR.

### Bulk RNA-sequencing

Murine aortic SMCs were isolated from embryos at E15.5 as described above and subjected to RNA extraction without culturing the cells. Human aortic SMCs in culture were treated with Scr RNA or siELN as described above and subjected to RNA extraction 72 hours after siRNA transfection. RNA was then sequenced using Illumina HiSeq 100 bp by paired-end sequencing at the Yale Center for Genome Analysis. Data analyses were performed using Partek Flow software (San Diego, CA, USA).

### qRT-PCR

The entire aorta from the root to the iliac arteries or the DA was dissected and homogenized with tissue homogenizer (Wheaton) in lysis buffer (ThermoFisher, 466001). RNA was isolated from tissue lysates as well as from cultured human and murine aortic or DA SMCs with the PureLink RNA Mini Kit (ThermoFisher, 12183018). Isolated RNA was reverse transcribed with the iScript cDNA Synthesis Kit (Bio-Rad). cDNA was subjected to qRT-PCR on a CFX96 Real-Time System (Bio-Rad) using SsoFast Eva-Green supermix (Bio-Rad) and primers as shown in Table S2. mRNA levels were normalized to 18S rRNA or Gapdh.

### Western blot

Aortas and DAs were mechanically lysed in RIPA buffer (ThermoFisher, 89901) with protease and phosphatase inhibitor cocktails (ThermoFisher, 1861281) on ice with a glass pestle tissue homogenizer (Pyrex). Cultured human or murine aortic SMCs were collected in RIPA buffer and vortexed every 10 minutes for 1 hour on ice. Lysates were then centrifuged at 13,000g, 4°C for 5 minutes, and supernatants were collected. BCA assay (ThermoFisher, 23225) was used to determine the protein concentration. Lysates were prepared in 4× Laemmli sample buffer (Bio-Rad) at 95°C for 5 minutes. Protein samples with Laemmli buffer were resolved by 4-20% SDS-PAGE, transferred to Immobilon PVDF membranes (Millipore), blocked with 5% nonfat dry milk or bovine serum albumin in TBS with tween 20 (TBS-T), washed in TBS-T, and incubated with primary antibodies diluted in blocking solution overnight at 4°C. On the next day, membranes were washed in TBS-T, incubated with HRP-conjugated secondary antibodies (Dako), washed in TBS-T again, and developed with Supersignal West Femto Maximum Sensitivity Substrate (ThermoFisher) on the G:BOX imaging system (Syngene). Western blot analysis used primary antibodies raised in rabbits and targeting SPHK1 (Proteintech, 10670-1-AP; 1:1000), EGR1 (Cell signaling, 4154; 1:1000), S1PR1 (Proteintech, 55133-1-AP, 1:1000), β-actin (Abcam, ab8227; 1:1000), human ELN (Elastin Products Company, PR533; 1:500), and ELN (raised against exons 6–17 of recombinant mouse tropoelastin; 1:500) (*12, 14, 67*).

### Detection of *Sphk1* deletion efficiency

To elucidate *Sphk1* deletion efficiency in SMCs, pregnant dams bearing *Sphk1(flox/flox)* embryos with no Cre or *Acta2-CreER^T2^* were injected daily with tamoxifen from E13.5-17.5, and pups were collected at P0.5 (*12*). From isolated aortic SMCs, RNA was purified for qRT-PCR, and protein was prepared for Western blot as described above. For genotyping, genomic DNA was extracted from the aortic media by incubation with 50 mM NaOH at 95°C for 1 hour, and then 100 mM Tris-HCl pH 7.5 was added to neutralize the digested samples. After centrifugation at 13,000g for 5 minutes, supernatants were used for genotyping PCR with primers as shown in Table S3.

As in our prior studies (*12, 14*), to determine gene deletion efficiency in ECs, pregnant dams bearing *Sphk1(flox/flox)* embryos with or without *Cdh5-CreER^T2^* were injected daily with tamoxifen from E13.5 to E17.5, and pups were collected at P0.5. Dissected lungs were incubated in 5 ml dissociation enzyme (Miltenyi, 130-095-927) at 37°C for 20 minutes (*12, 14*). Samples were then processed with the gentleMACS Dissociator (Miltenyi, 130-093-235). The lung cell suspension was filtered with 70 μm strainer, and the enzymatic reaction was stopped with 1 ml FBS. The resulting cell suspension was centrifuged at 500 rpm for 5 minutes, and the cell pellet was resuspended in M199 medium. Subsequently, magnetic beads (Invitrogen, 11035), which were pre-coated with anti-mouse-CD31 antibody (BD Pharmingen, 553370), were added to the cell suspensions and together were incubated on a rotator for 20 minutes. ECs bound to magnetic beads were collected with a magnet stand and used for PCR, qRT-PCR, and Western blot (*12, 14*).

### SMC proliferation assay

Aortic SMCs isolated from pups at P0.5 were cultured with EdU (10 µM) from the Click-iT EdU Alexa Fluor 647 Imaging Kit (ThermoFisher, C10338) for 8 hours. Cells were then fixed with 4% PFA for 30 min, permeabilized in 0.5% Triton X-100 in PBS and stained for EdU and nuclei (Hoechst).

### SMC migration assay

Aortic SMCs isolated from pups at P0.5 by enzyme mixture were transfected with Scr RNA or siSphk1 as described above. SMCs were scratched with pipet tips 72 hours after siRNA treatment and washed with M199 medium. The cell coverage of the scratched area was measured immediately after the scratch and 12, 24, 36, and 48 hours later.

### mRNA decay assay

mRNA decay rates were assessed using transcription inhibitor (5,6-Dichlorobenzimidazole 1-β-D-ribofuranoside [DRB]) (Sigma-Aldrich, D1916). Isolated murine aortic SMCs were seeded on plates. After starvation with 0.5% FBS for 24 hours, cells were treated with DRB (60 μM). Samples were collected with lysis buffer (PureLink RNA Mini Kit, ThermoFisher) at 0, 15, 30 minutes, 1, 3, 6, 12, and 24 hours after DRB treatment, and Sphk1 mRNA levels were analyzed by qRT-PCR.

### Decellularization

Murine aortic SMCs were cultured with 10% FBS supplemented medium for 10 days without changing the medium to facilitate elastic fiber assembly (*34*). Decellularization was performed by freeze-thaw cycles and detergent treatments (*12*). Briefly, murine aortic SMCs were placed in -80°C for 30 minutes and then washed with PBS at room temperature for 30 minutes. This freeze-thaw cycle was repeated three times. Cells were then treated with 0.5% NP-40 in PBS for 30 minutes at room temperature and washed three times in PBS for 10 minutes.

CF633 hydrazide was used for elastin staining (Sigma-Aldrich, SCJ4600037; 1:3000) (*12, 66*). The resulting decellularized matrices were incubated at 37°C for 1 hour in SMC culture medium and then murine aortic SMCs were re-seeded at 90% confluency on these matrices and incubated for 48 hours. SMC lysates were collected in lysis buffer and subjected to qRT-PCR.

### Stiffness assay

Polydimethylsiloxane (PDMS) substrate A and B, liquid oligomeric base and crosslinker reagents (Sylgard184, Electron Microscopy Science, 24236-10) were used for preparing surfaces of varying stiffness (*36*). PDMS substrate A and B at different proportions (A:B = 5:1, 7.5:1, 10:1, 40:1, and 60:1) were mixed by vigorous stirring and then, 1 ml of this pre-polymer was poured onto a 6-well plate and cross-linked in an oven at 80 °C for 3 hours. The prepared PDMS surface was coated with fibronectin (10 μg/ml, purified from bovine plasma (*68*)) at 4 °C overnight and sterilized using ultraviolet light for 30 minutes before use.

### Vascular casting

After euthanizing pups with isoflurane, the chest wall was opened, and left lower extremity was removed. PBS was injected in the left ventricle and blood was drained through the left iliac artery. Yellow latex (Ward’s Science, 470024-616) was then injected through the left ventricle (*14*). Mice were kept moist at 4°C for 3-4 hours to facilitate setting of the injected latex and subsequently, fixed in 4% PFA at 4 °C overnight, followed by dissection to reveal the aorta (*14*).

### Mean linear intercept

Mean linear intercept was used to assess lung emphysema (*14*). Lung cryosections were subjected to H&E staining, and images were collected using a 10x objective for 3 sections per lung. A grid with 5 lines (500 μm in length per line) was placed over each image. The value of one intercept was defined as the linear length between epithelium of two adjacent alveoli. For each line, the values of all intercepts were quantified. A larger value represents more simplified alveoli.

### Imaging

Fluorescence images of aortic sections were acquired with a confocal microscope (PerkinElmer UltraView Vox Spinning Disc). Brightfield images of H&E staining were captured using a BX63 microscope (Olympus). Adobe Photoshop (version 25.9.0) was used for image processing.

### Statistical analysis

Two-tailed Student’s *t* test and multifactor ANOVA with Tukey’s *post hoc* test were used to analyze data using GraphPad Prism (version 9.4.1). Statistical significance threshold was set at a *P* value of less than 0.05. All data are presented as mean ± SD.

## Acknowledgments

We thank Dr. David Montefusco at Virginia Commonwealth University for providing *Sphk1(flox/flox)* mice, Dr. Martin Schwartz at Yale University School of Medicine for providing fibronectin, and Mr. Rolando Garcia-Milian at Yale University for supporting bulk RNA-seq analyses. We also thank Greif laboratory members for their input.

## Author contributions

J.S., J.M.D., and D.M.G. conceived of and designed experiments. J.S., J.M.D., and E.G-V. performed them. G.T., R.K.R, and Z.U. provided human aortic samples. J.M.D., I.K., and T.H. provided input for study direction. T.H. provided *S1pr1(knock-in/knock-in), H2B-GFP* mice. J.S. and D.M.G. analyzed the results, prepared the figures, and wrote the manuscript. All authors reviewed and provided input on the manuscript.

## List of Supplementary Materials

- Figures S1 to S15

- Supplementary Figures Legends

- Tables S1-3

---

**Figure S1.**
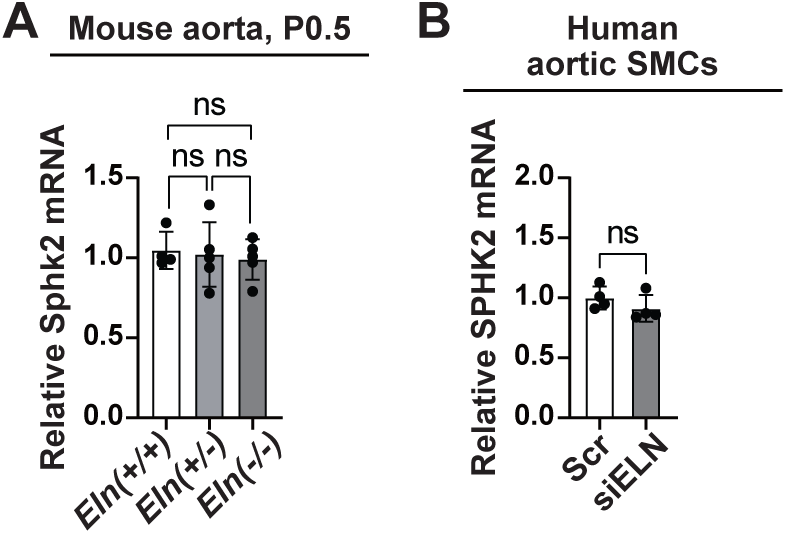
Elastin insufficiency does not change expression of SPHK2 in mouse aorta and human aortic SMCs. **(A, B)** Aortic lysates from *Eln(+/+)*, *Eln(+/-)* or *Eln(-/-)* mice at P0.5 (**A**) or from human aortic SMCs treated with scrambled (Scr) or siELN (**B**) were subjected to qRT-PCR for SPHK2. n=4-5. ns, not significant by multifactor ANOVA with Tukey’s *post hoc* test (**A**) or Student’s *t* test (**B**).

**Figure S2.**
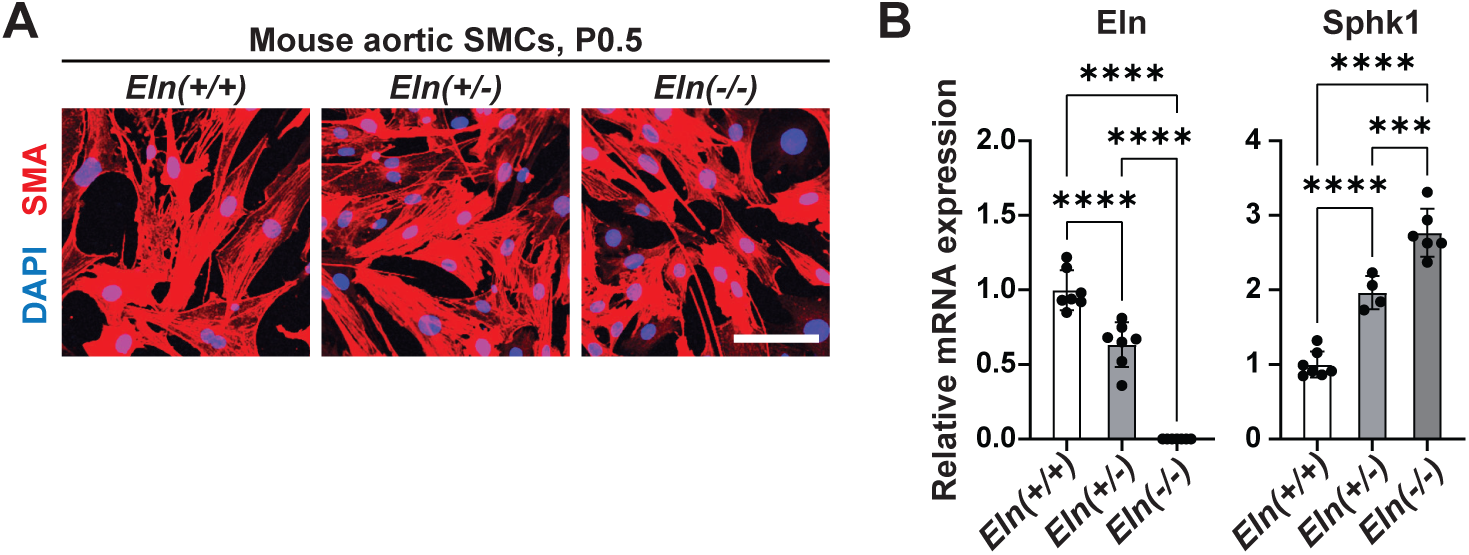
Reduced *Eln* gene dosage increases Sphk1 transcript in cultured murine aortic SMCs. Aortic SMCs were isolated from *Eln(+/+)*, *Eln(+/-)* or *Eln(-/-)* pups at P0.5. **(A)** Immunostaining for SMA and nuclei (DAPI). Scale bar, 100 μm. **(B)** Lysates from SMCs were subjected to qRT-PCR for Eln or Sphk1. n=4-7. \*\*\**P* < 0.001, \*\*\*\**P* < 0.0001 by multifactor ANOVA with Tukey’s *post hoc* test.

**Figure S3.**
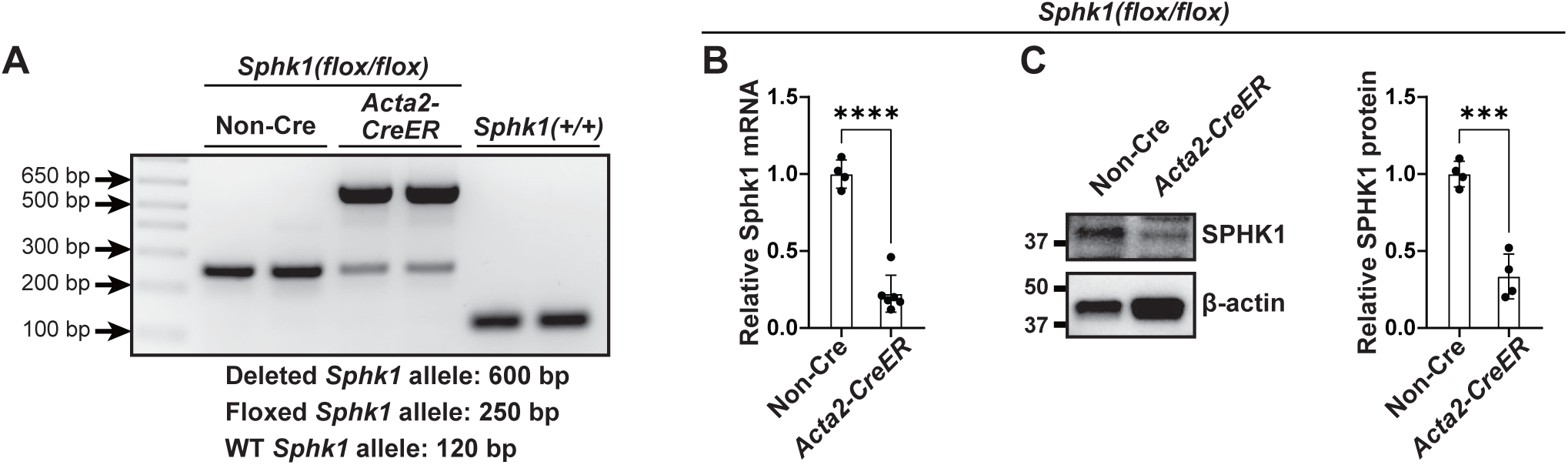
*Sphk1* deletion efficiency in aortic SMCs of *Acta2-CreER^T2^*, *Sphk1(flox/flox)* mice. Dams pregnant with *Sphk1(flox/flox)* embryos also carrying *Acta2-CreER^T2^* or no Cre were injected daily with tamoxifen from E13.5 to E17.5. Aortas were harvested from pups at P0.5 and subjected to lysis after adventitia and EC removal. **(A)** Extracted genomic DNA was subjected to PCR with primers flanking *Sphk1*. The 120, 250, and 600 PCR products represent the wild-type, floxed, and deleted *Sphk1* alleles, respectively. **(B)** Isolated RNA was reverse transcribed and subjected to qRT-PCR. Histogram represents Sphk1 transcript levels relative to Gapdh. n=4-6. **(C)** Western blot was performed on aortic lysates. Densitometry of protein bands relative to β-actin is shown. n=4 mice. \*\*\**P* < 0.001, \*\*\*\**P* < 0.0001 by Student’s *t* test (**B, C**).

**Figure S4.**
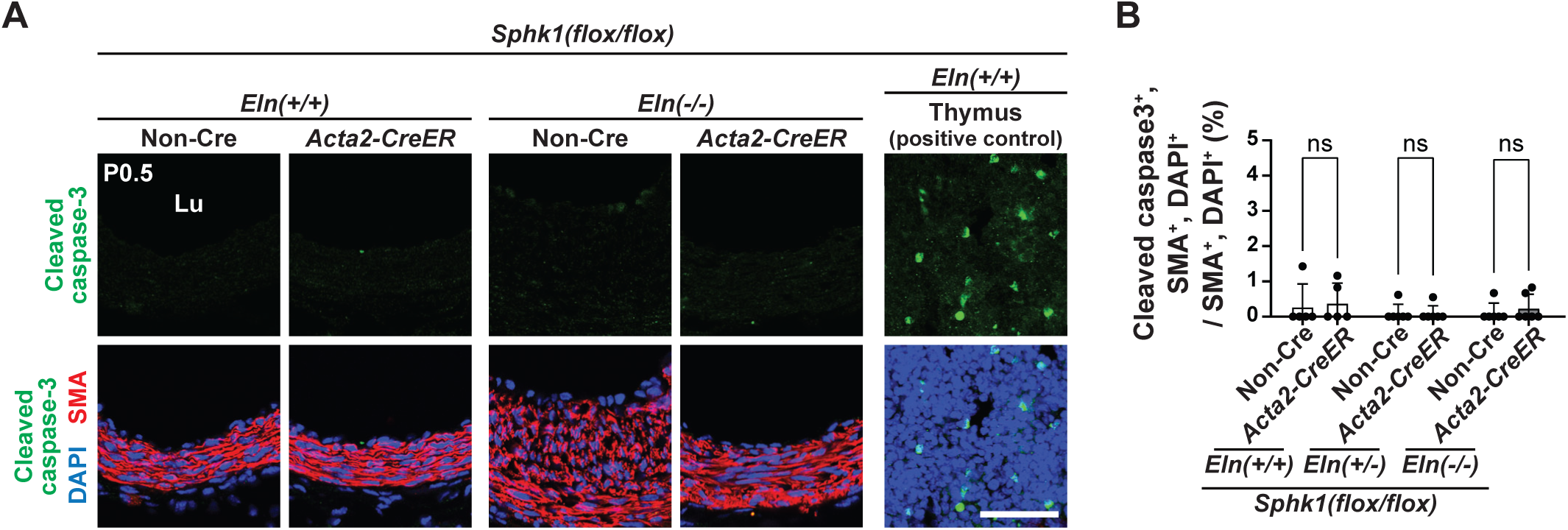
*Sphk1* deletion in SMCs does not induce apoptosis in aortic SMCs. **(A)** Pregnant dams were injected with tamoxifen daily from E13.5 to E17.5, and pups were collected at P0.5. Transverse aortic sections were stained for markers of apoptosis (cleaved caspase-3), SMCs (SMA), and nuclei (DAPI). Section of the thymus from *Eln(+/+),* non-Cre pup at P0.5 is a positive control for cleaved caspase-3 staining. Scale bar, 50 μm. **(B)** Percent of SMCs that are cleaved caspase-3^+^ from sections represented in **A**. n=5-6 mice. ns, not significant by multifactor ANOVA with Tukey’s *post hoc* test.

**Figure S5.**
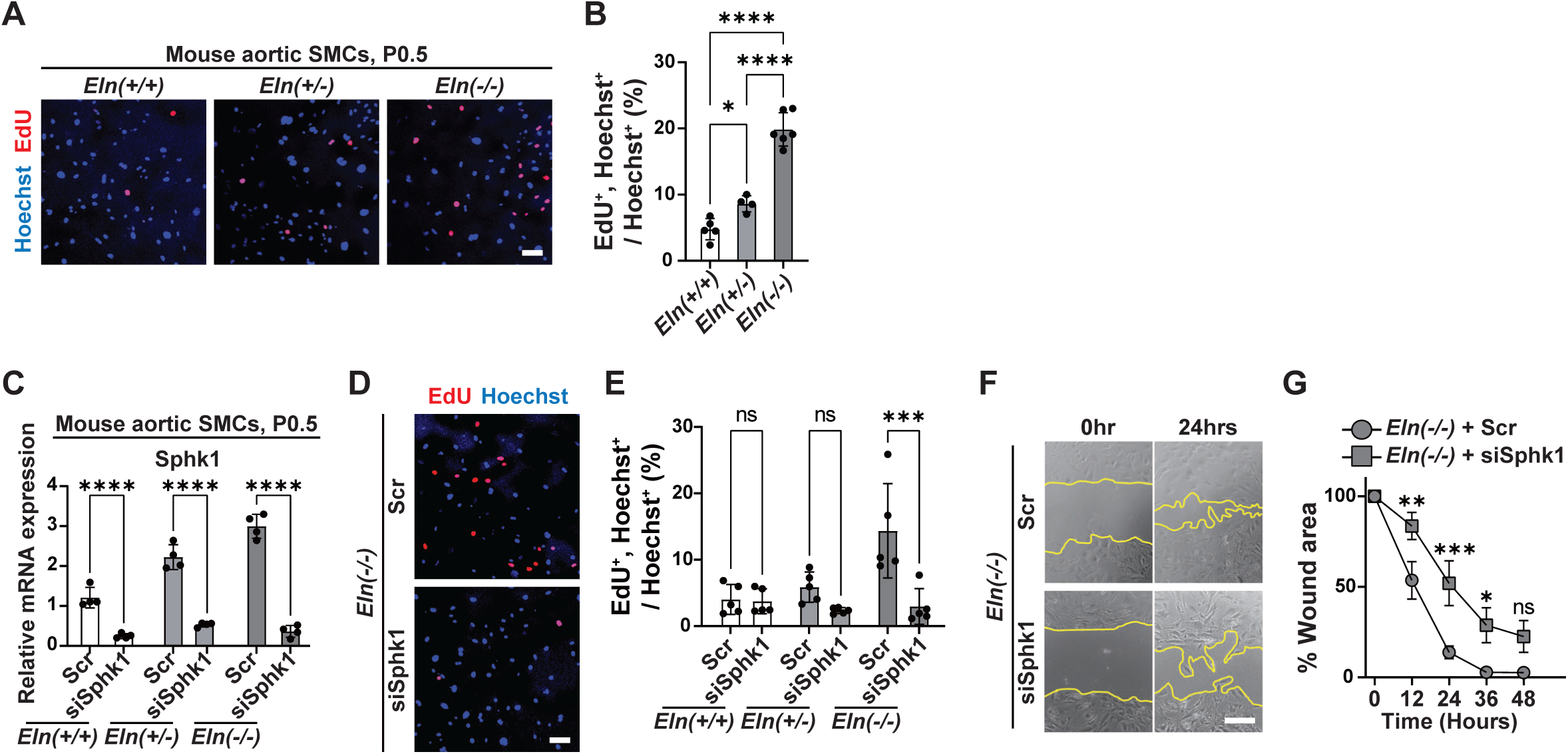
Inhibition of SPHK1 attenuates SMC proliferation and migration with elastin deficiency. Aortic SMCs were isolated from *Eln(+/+)*, *Eln(+/-)* or *Eln(-/-)* pups at P0.5. **(A, B)** SMCs were cultured with EdU for 8 hours and stained for EdU and nuclei (Hoechst). n=4-6. **(C)** SMCs were treated with scrambled (Scr) RNA or siSphk1 and then subjected to qRT-PCR. n=4. **(D, E)** SMCs treated with Scr RNA or siSphk1 were cultured with EdU for 8 hours and then stained for EdU and nuclei (Hoechst). n=5. **(F, G)** SMCs were treated with siRNA and then subjected to scratch assay. Percent of wound area relative to time 0 was quantified. n=5-6. ns, not significant, \**P* < 0.05, \*\**P* < 0.01, \*\*\**P* < 0.001, \*\*\*\**P* < 0.0001 by multifactor ANOVA with Tukey’s *post hoc* test (**B, C, E**) and Student’s *t* test (**G**). Scale bars, 100 μm (**A, D, F**).

**Figure S6.**
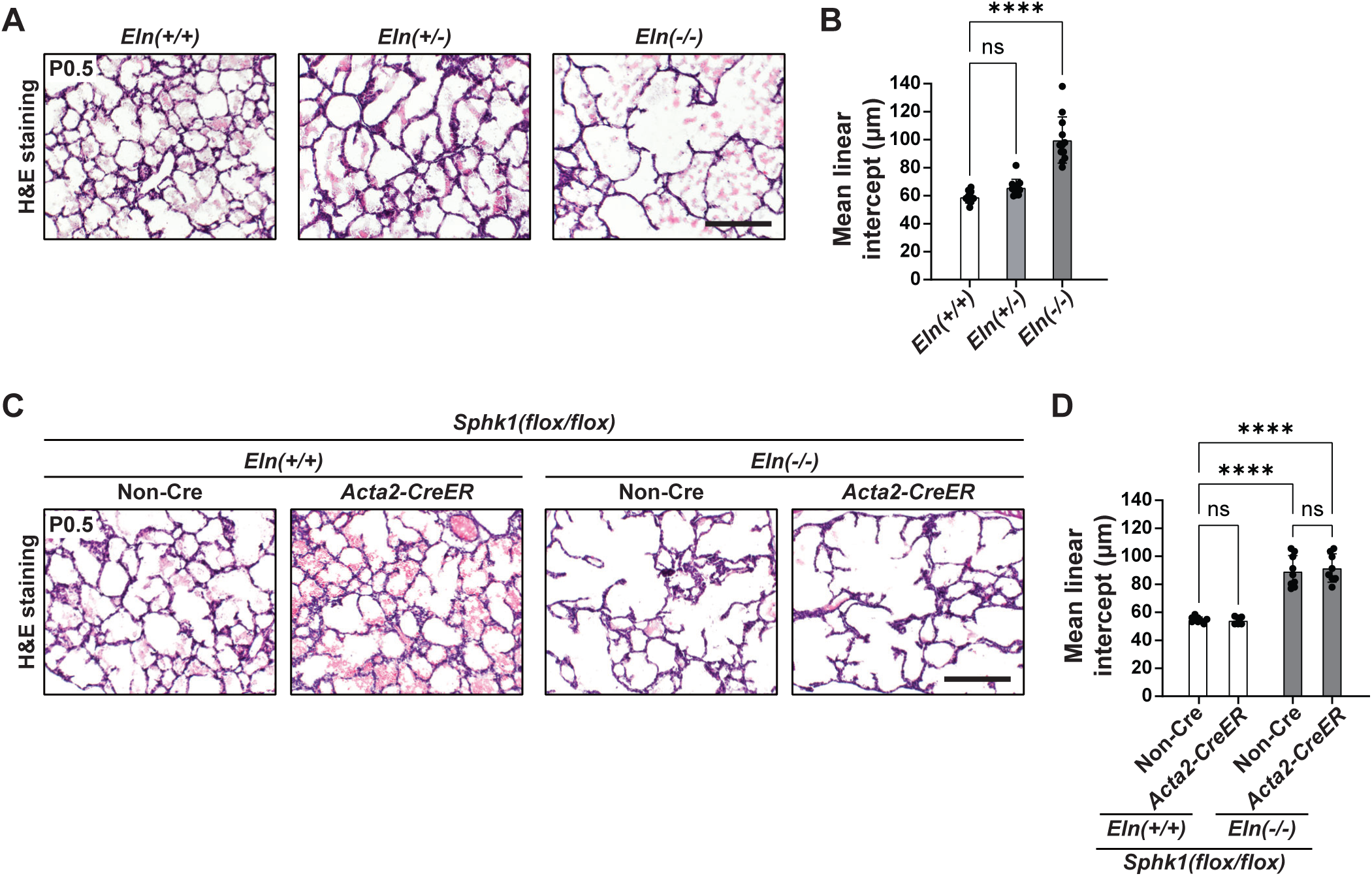
*Sphk1* deletion in SMCs does not improve emphysema in *Eln(-/-)* mice. **(A, C)** Transverse lung sections from mice of indicated genotypes at P0.5 were subjected to H&E staining. Of note in **C**, pregnant dams were injected with tamoxifen daily from E13.5 to E17.5 . **(B, D)** Histograms represent mean linear intercept from sections as shown in **A** and **C**. n=6-12 mice. ns, not significant, \*\*\*\**P* < 0.0001 by multifactor ANOVA with Tukey’s *post hoc* test. Scale bars, 200 μm (**A, C**).

**Figure S7.**
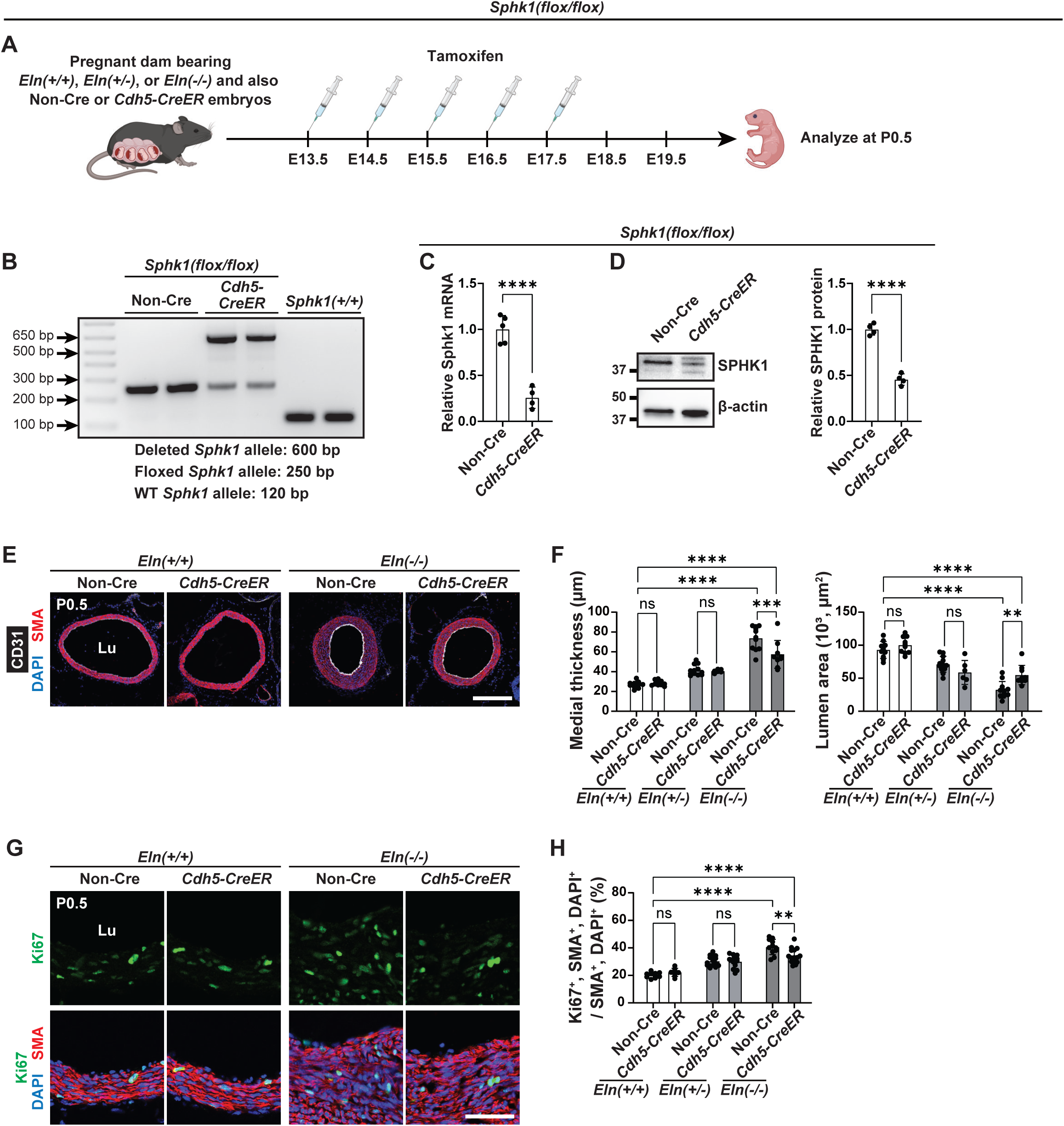
*Sphk1* deletion with *Cdh5-CreER^T2^* modestly attenuates elastin aortopathy. **(A)** Experimental strategy for (**B-H**). Pregnant dams were injected with tamoxifen daily from E13.5 to E17.5, and pups were collected at P0.5. **(B)** Extracted genomic DNA was subjected to PCR with primers flanking *Sphk1*. The 120, 250, and 600 bp PCR products represent the wild-type, floxed, and deleted *Sphk1* alleles, respectively. **(C)** Isolated RNA was reverse transcribed and subjected to qRT-PCR. Histograms represent Sphk1 transcript levels relative to Gapdh. n=4-5. **(D)** Western blot was performed on lysates of ECs isolated from the aorta. Densitometry of protein bands relative to β-actin is shown on right. n=4 mice. **(E)** Transverse aortic sections of indicated genotypes were stained for SMA, CD31, and nuclei (DAPI). **(F)** Histograms represent aortic medial thickness and lumen area from sections as shown in **E**. n=6-12 mice. **(G)** Transverse aortic sections were stained for proliferation marker Ki67, SMA, and nuclei (DAPI). **(H)** Histograms represent percent of SMCs that are Ki67^+^ as in **G**. n=7-15 mice. ns, not significant, \*\**P* < 0.01, \*\*\*\**P* < 0.0001 by Student’s *t* test (**C, D**) and multifactor ANOVA with Tukey’s *post hoc* test (**F, H**). Scale bars, 200 μm (**E**), 50 μm (**G**).

**Figure S8.**
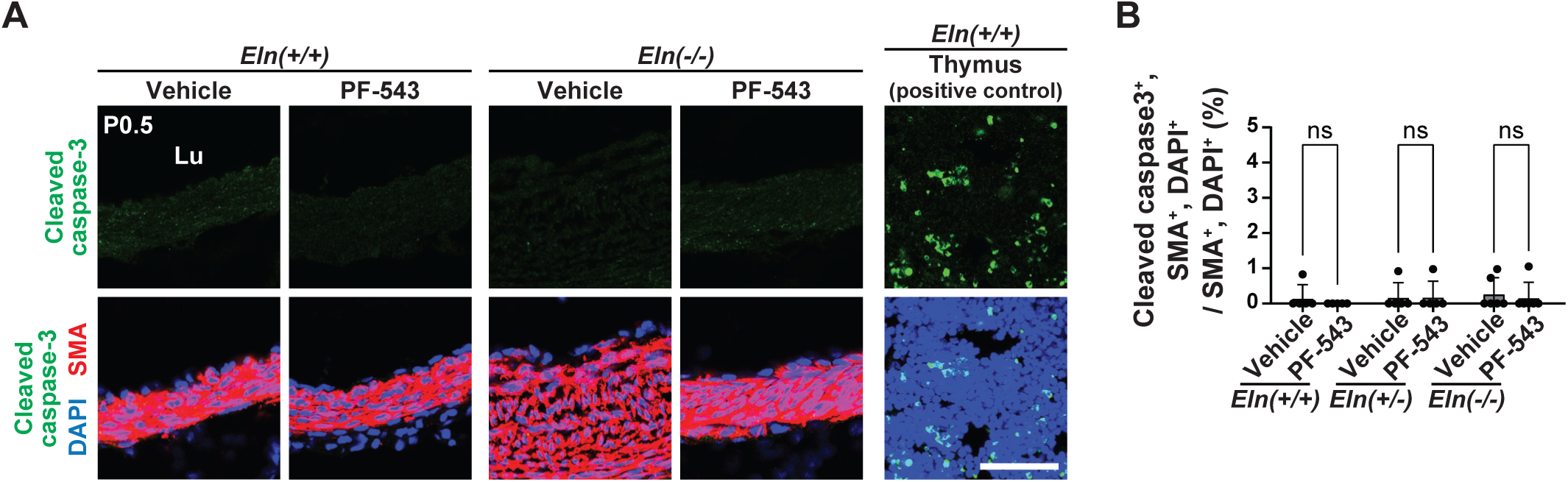
SPHK1 selective inhibitor does not induce apoptosis in aortic SMCs. **(A)** Pregnant dams were injected with vehicle or SPHK1 inhibitor (PF-543) daily from E13.5 to E19.5, and pups were collected at P0.5. Transverse aortic sections of *Eln(+/+)* and *Eln(-/-)* pups were stained for cleaved caspase-3, SMA, and nuclei (DAPI). Section of thymus from *Eln(+/+)* pup injected with vehicle at P0.5 was a positive control for cleaved caspase-3 staining. **(B)** Percent of SMCs that are cleaved caspase-3^+^ from sections represented in **A**. n=5-6 mice. ns, not significant by multifactor ANOVA with Tukey’s *post hoc* test. Scale bar, 50 μm.

**Figure S9.**
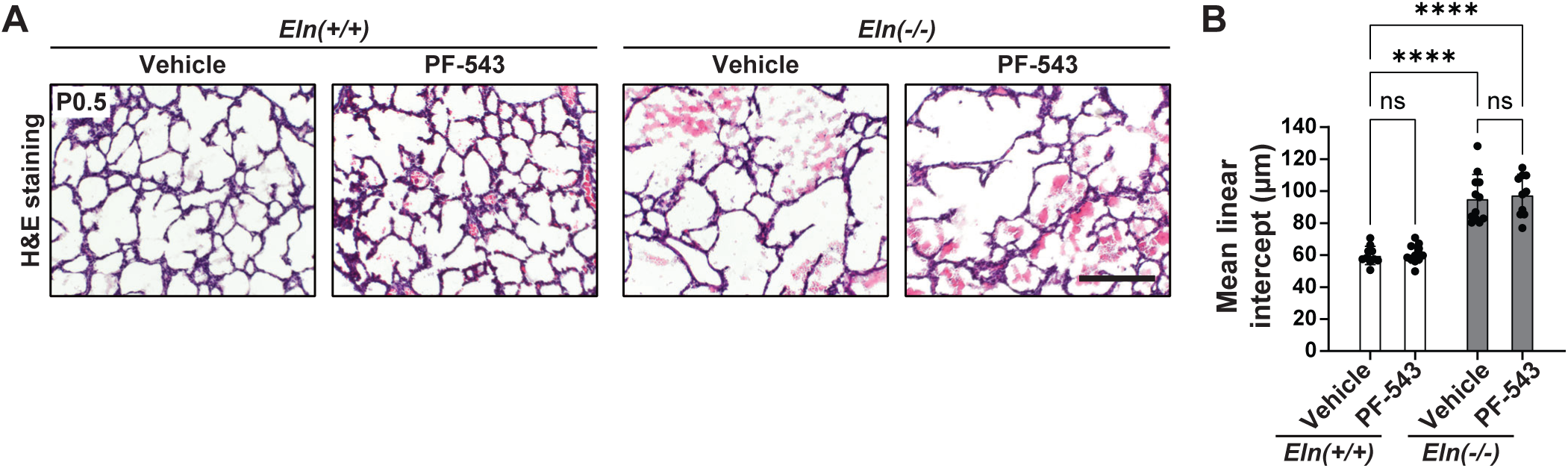
SPHK1 selective inhibitor does not improve emphysema in *Eln(-/-)* mice. Pregnant dams were injected with vehicle or SPHK1 inhibitor (PF-543) daily from E13.5 to E19.5, and pups were collected at P0.5. Lung transverse sections from *Eln(+/+)* and *Eln(-/-)* pups were subjected to H&E staining. Scale bar, 200 μm. **(B)** Histograms represent mean linear intercept from sections as shown in **A**. n=10-12 mice. ns, not significant, \*\*\*\**P* < 0.0001 by multifactor ANOVA with Tukey’s *post hoc* test.

**Figure S10.**
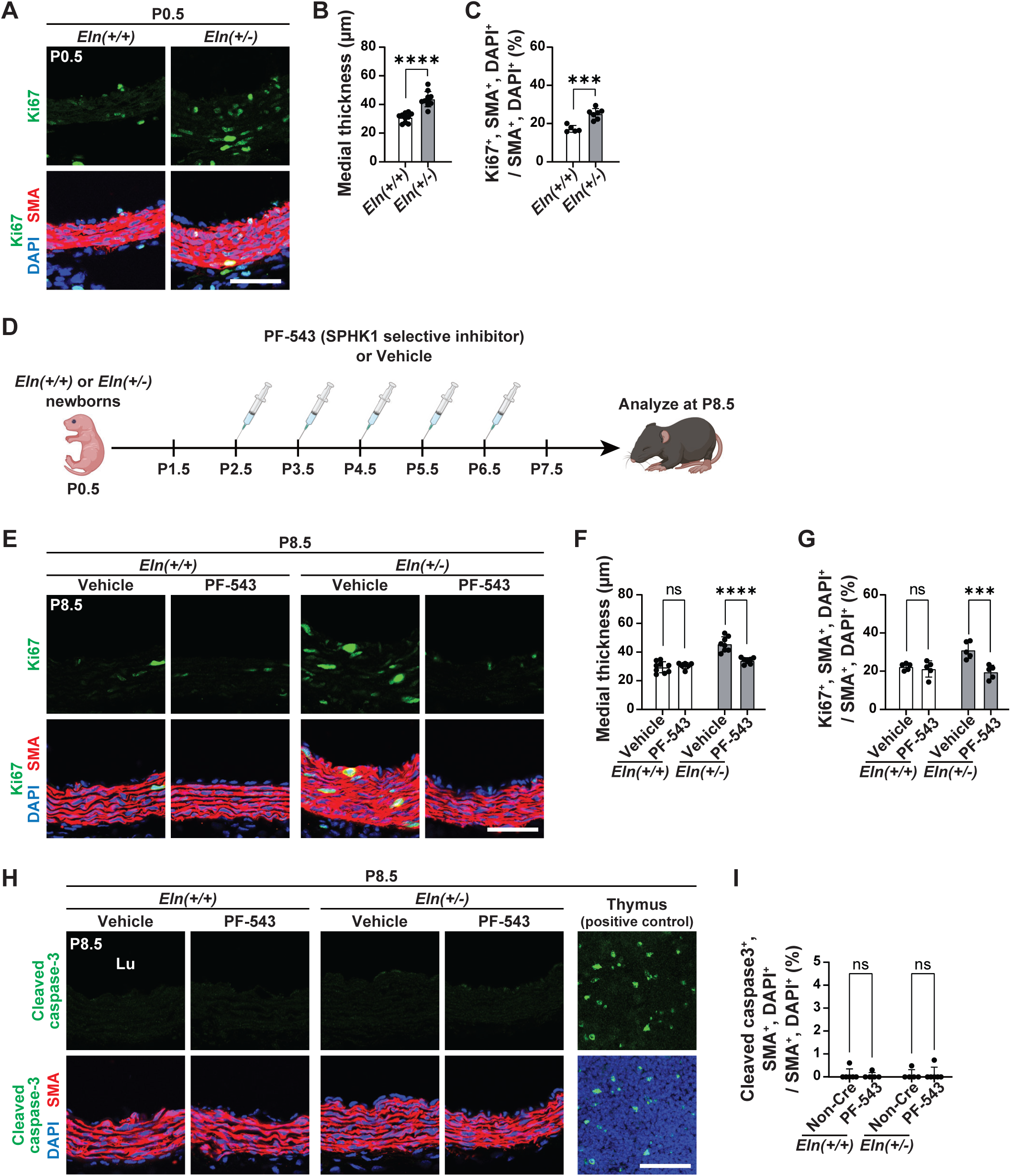
SPHK1 selective inhibitor mitigates established aortic muscularization in *Eln(+/-)* mice. **(A)** Transverse aortic sections from *Eln(+/+)* or *Eln(+/-)* pups at P0.5 were stained for Ki67, SMA, and nuclei (DAPI). **(B)** Histograms represent aortic medial wall thickness from sections as shown in **A**. n=10 mice. **(C)** Percent of SMCs that are Ki67^+^ from sections represented in **A**. n=5-7 mice. **(D)** Experimental strategy for (**E-I**). *Eln(+/+)* or *Eln(+/-)* pups were injected daily with PF-543 (20 mg/kg body weight) or vehicle from P2.5 to P6.5 and analyzed at P8.5. **(E, H)** Transverse aortic sections were stained for SMA, nuclei (DAPI) and either Ki67 in **E** or cleaved caspase 3 in **H**. **(F)** Histograms represent aortic medial wall thickness from sections at P8.5 as shown in **E**. n=7-9 mice. **(G)** Percent of SMCs that are Ki67^+^ from sections represented in **E**. n=5 mice. **(I)** Percent of SMCs that are cleaved caspase3^+^ from sections represented in **H**. n=5-6 mice. ns, not significant, \*\*\**P* < 0.001, \*\*\*\**P* < 0.0001 by Student’s *t* test (**B, C**) and multifactor ANOVA with Tukey’s *post hoc* test (**F, G, I**). Scale bars, 50 μm (**A, E, H**).

**Figure S11.**
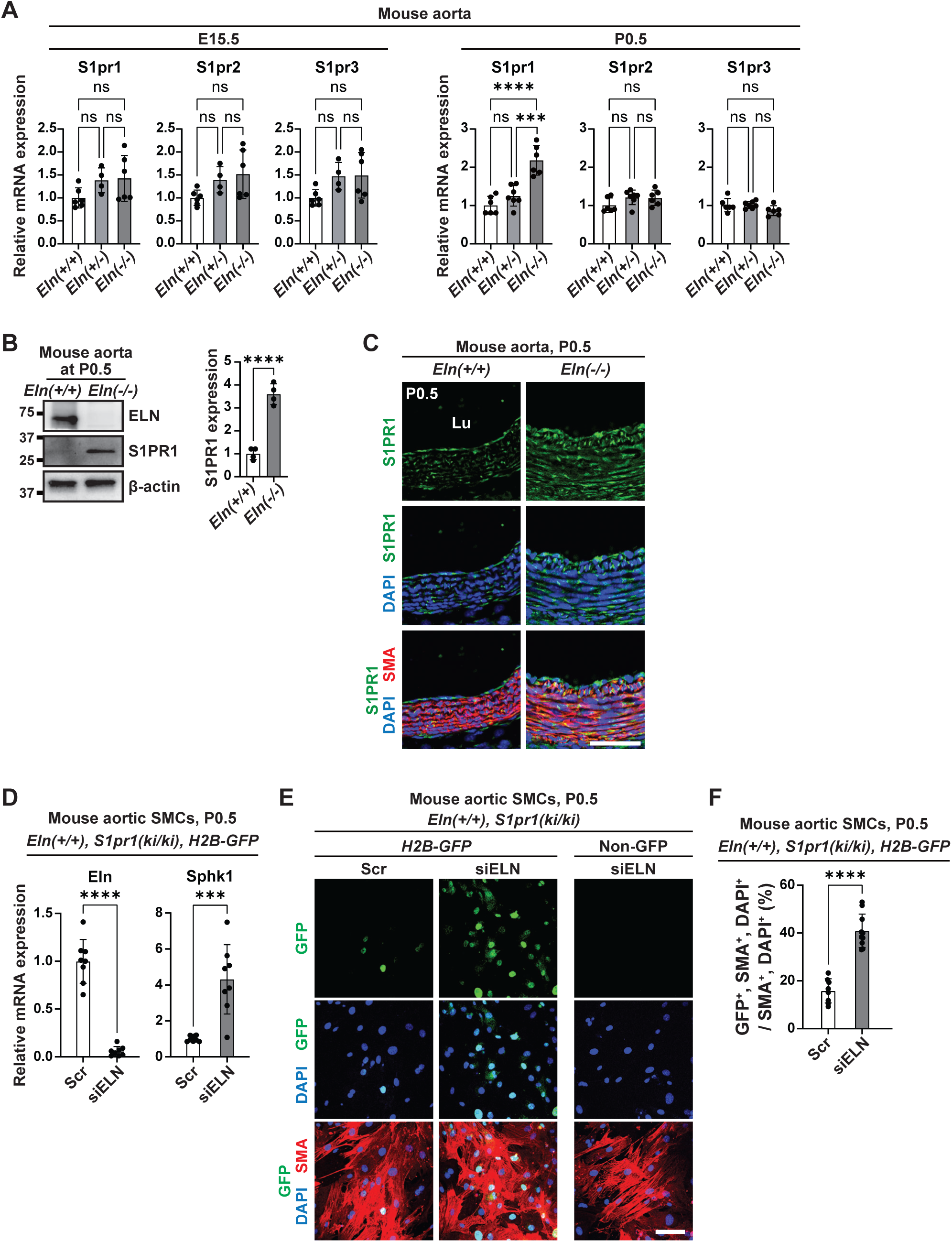
S1PR1 is activated by elastin insufficiency. **(A, B)** Lysates from aortas of *Eln(+/+)*, *Eln(+/-)* or *Eln(-/-)* mice at E15.5 and P0.5 were subjected to qRT-PCR for S1pr1, S1pr2 and S1pr3 (**A**) or Western blotting for S1PR1 (**B**). Densitometry of bands relative to β-actin is shown in **B**, right. n=4-7. **(C)** Ascending aortic sections of *Eln(+/+)* or *Eln(-/-)* mice at P0.5 stained for S1PR1, SMA, and nuclei (DAPI). n=4-6 mice. Lu, lumen. **(D-F)** Aortic SMCs isolated from *Eln(+/+)*, *S1pr1(knock-in/knock-in)*, *H2B-GFP* pups at P0.5 were treated with Scr RNA or siEln. **(D)** Lysates from SMCs treated with siRNA as indicated were subjected to qRT-PCR for Eln and Sphk1. n=8. **(E)** SMCs with indicated genotypes and treatments were stained for GFP (marker for S1PR1 activity), SMA, and nuclei (DAPI). **(F)** Histograms represent percent of SMCs that are GFP^+^ as in **E**. n=8-10. ns, not significant, \*\**P* < 0.01, \*\*\**P* < 0.001, \*\*\*\**P* < 0.0001 by multifactor ANOVA with Tukey’s *post hoc* test (**A**) or Student’s *t* test (**B, D, F**). Scale bars, 50 μm (**C**) and 100 μm (**E**).

**Figure S12.**
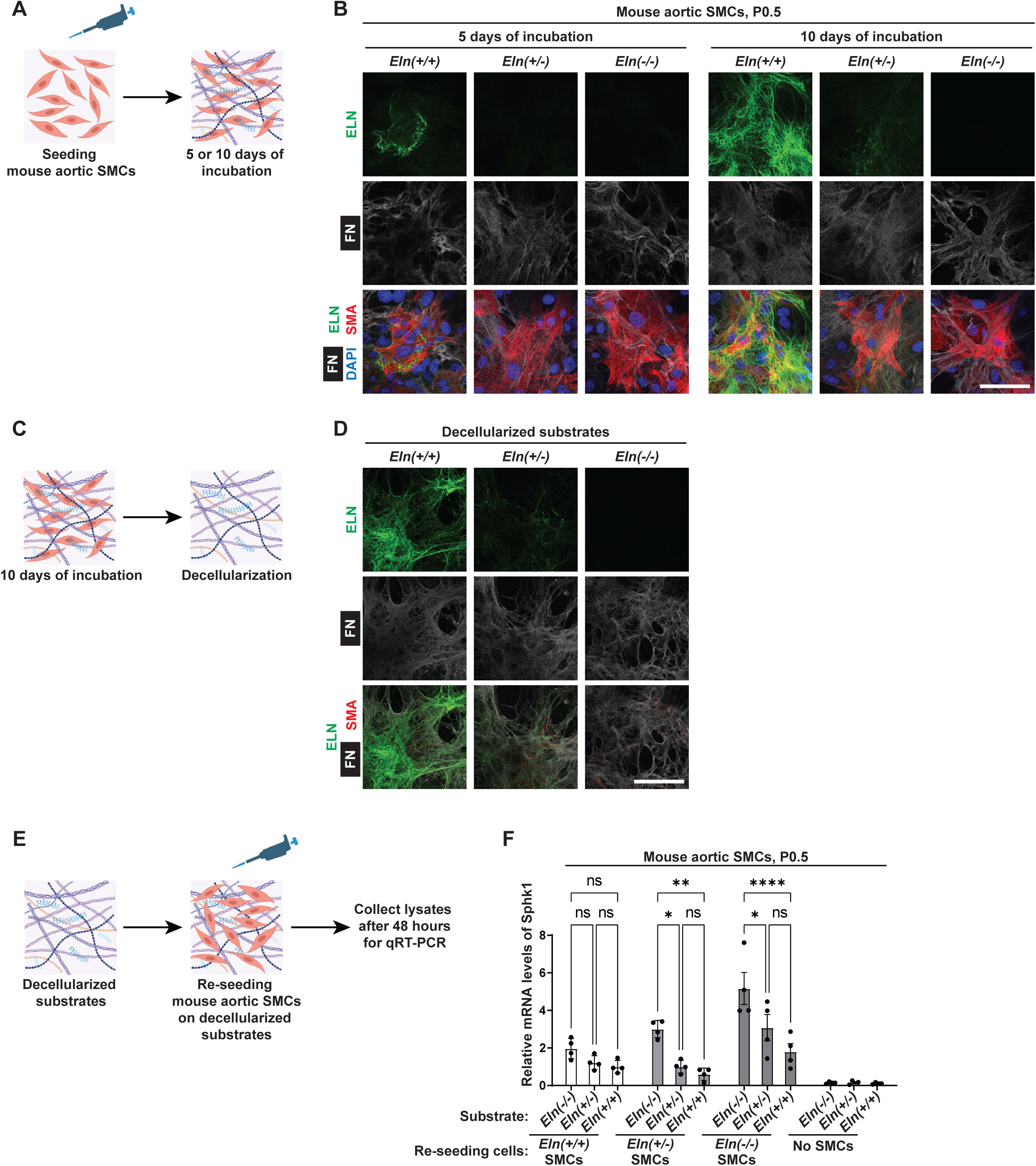
*Eln(+/+)* SMC-derived extracellular matrices attenuate Sphk1 expression in elastin mutant SMCs. **(A, B**) Isolated aortic SMCs from *Eln(+/+)*, *Eln(+/-)*, or *Eln(-/-)* pups at P0.5 were cultured for 5 or 10 days and subjected to staining for ELN, fibronectin (FN), SMA, and nuclei (DAPI). **(C, D)** After 10 days of incubation, cells were decellularized and immunostained for ELN, FN, and SMA. **(E)** After decellularization, aortic SMCs isolated from *Eln(+/+)*, *Eln(+/-)*, or *Eln(-/-)* pups at P0.5 were re-seeded on the resultant matrices. These samples were subjected to qRT-PCR after incubation for 48 hours. **(F)** Lysates from indicated SMCs on each substrate were subjected to qRT-PCR for Sphk1. n=4. ns, not significant, \**P* < 0.05, \*\**P* < 0.01, \*\*\*\**P* < 0.0001 by multifactor ANOVA with Tukey’s *post hoc* test. Scale bars, 100 μm (**B, D**).

**Figure S13.**
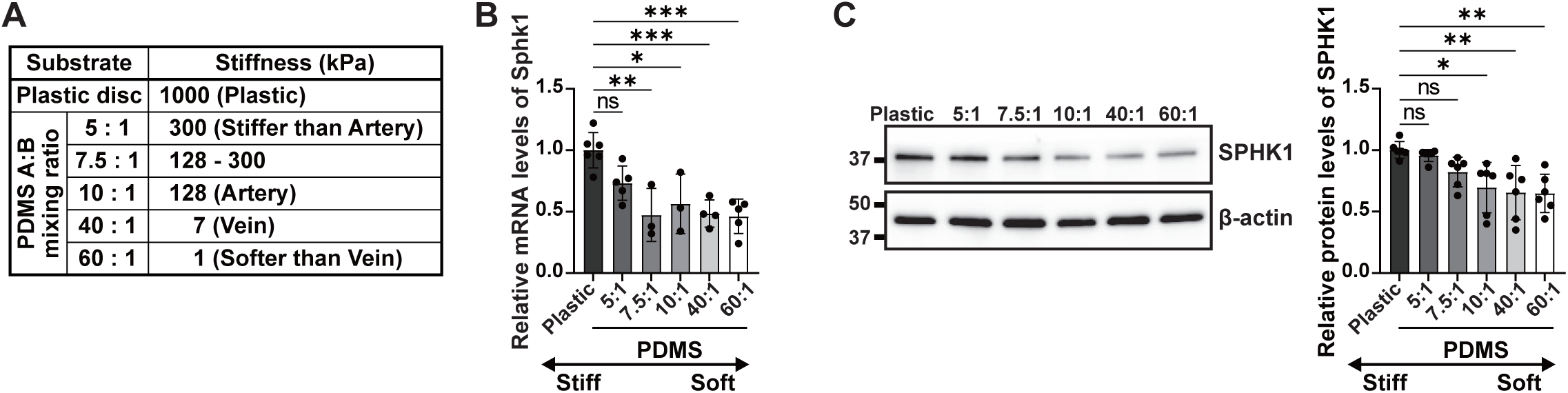
SPHK1 expression is upregulated by stiff substrates. **(A)** PDMS substrate A and B are liquid oligomeric base and crosslinker reagents (Sylgard184) with estimated stiffness. **(B, C)** Aortic SMCs isolated from *Eln(-/-)* pups at P0.5 were cultured on PDMS substrates for 72 hours and then subjected to qRT-PCR and Western blotting for SPHK1. Densitometry of bands relative to β-actin and normalized to this ratio on plastic is shown in **C**, right. n=3-6. ns, not significant, \**P* < 0.05, \*\**P* < 0.01, \*\*\**P* < 0.001 by multifactor ANOVA with Tukey’s *post hoc* test.

**Figure S14.**
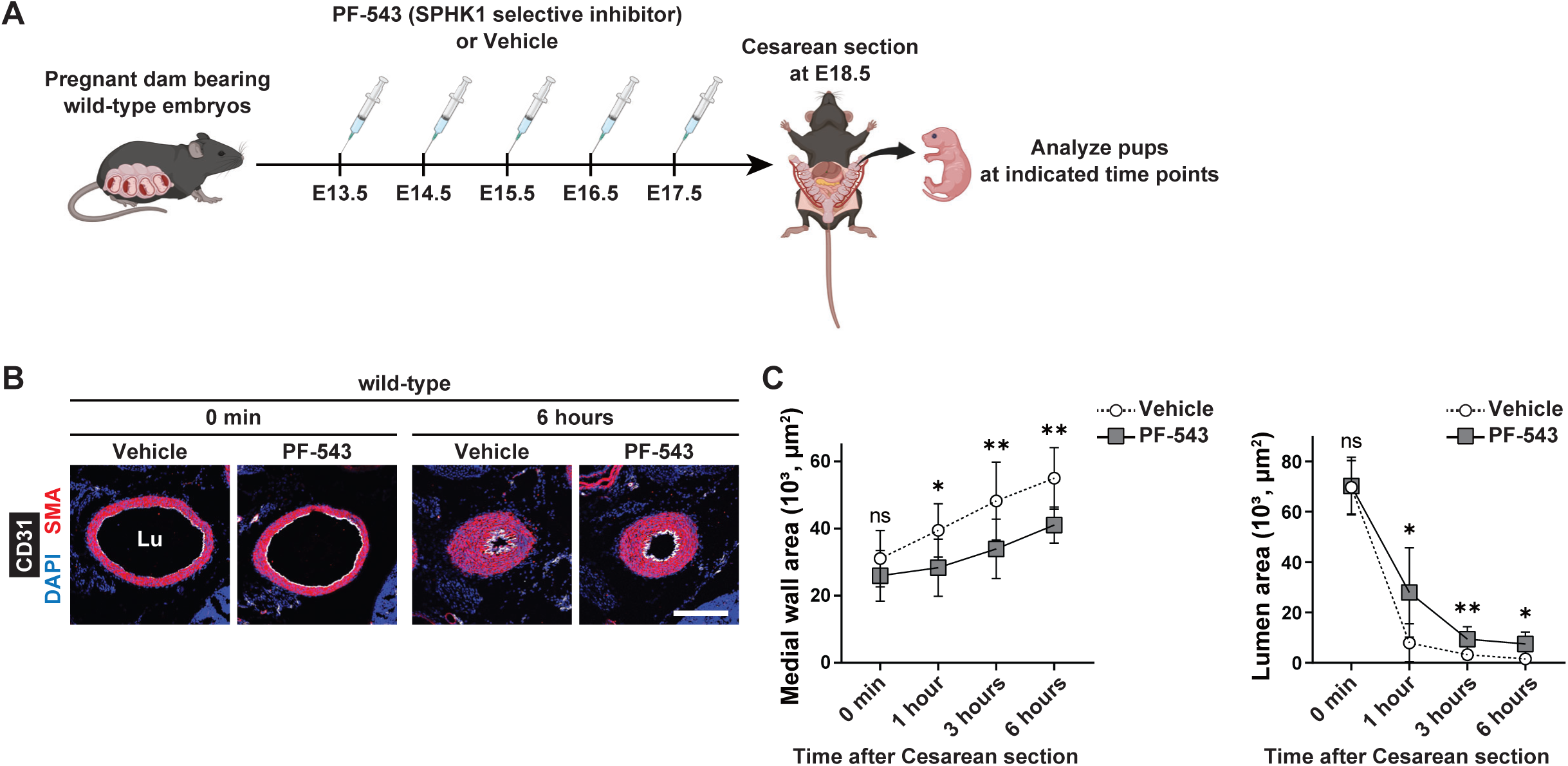
SPHK1 selective inhibitor induces persistent DA patency after birth. **(A)** Experimental strategy for (**B, C**). Pregnant dams bearing wild-type embryos were injected daily from E13.5-17.5 with SPHK1 inhibitor (PF-543, 20 mg/kg) or vehicle and subjected to cesarean section at E18.5. Neonates were kept alive and warm on a heating pad and harvested at postnatal ages up to 6 hours as indicated. **(B)** Representative images of transverse sections of DA at postnatal 6 hours stained for SMA, CD31, and nuclei (DAPI). Scale bar, 200 μm. **(C)** Medial wall and lumen area of DA at indicated time points after cesarean section. n=8-12 mice. ns, not significant, \**P* < 0.05, \*\**P* < 0.01 by Student’s *t* test.

**Figure S15.**
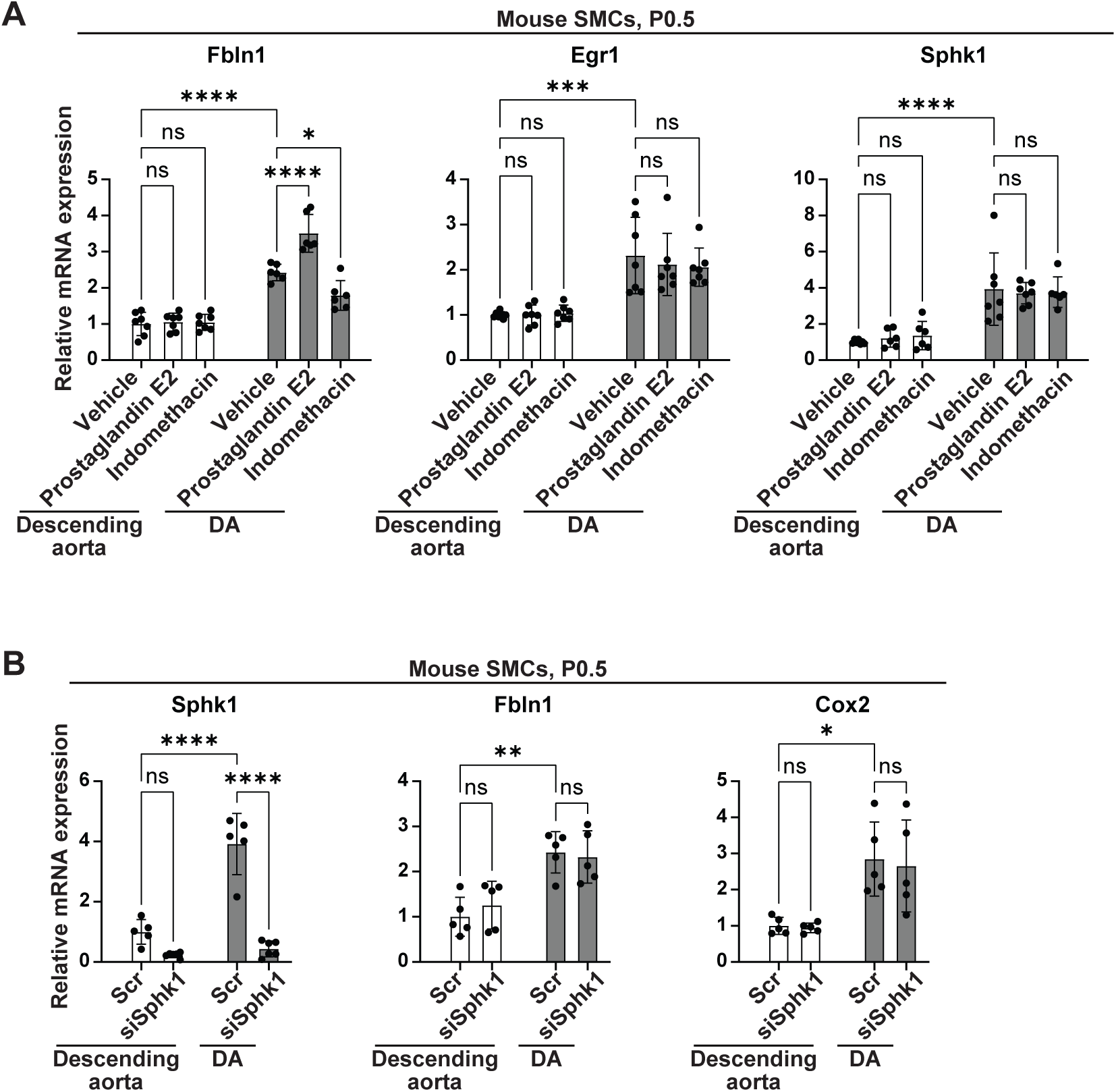
SPHK1 signaling is independent of prostaglandin E (PGE) pathway in DA SMCs. Descending aortic and DA SMCs were isolated from wild-type mice at P0.5. **(A)** SMCs were treated with either vehicle (DMSO), PGE2 (10^-6^ M), or indomethacin (10-5 M), and lysates were subjected to qRT-PCR for fibulin 1 (Fbln1), Egr1, and Sphk1. n=6-7. **(B)** SMCs were treated with Scr RNA or siSphk1, and lysates were subjected to qRT-PCR for Sphk1, Fbln1, and cyclooxygenase 2 (Cox2). n=5-6. ns, not significant, \**P* < 0.05, \*\**P* < 0.01, \*\*\**P* < 0.001, \*\*\*\**P* < 0.0001 by multifactor ANOVA with Tukey’s *post hoc* test.

**Table S1.** Demographic information for human WBS patients and controls.

| Age-matched control aorta |  |  | WBS aorta |  |  |
| --- | --- | --- | --- | --- | --- |
| Sample | Sex | Age | Sample | Sex | Age |
| Control | M | 59 | WBS | unknown | 42 |
| Control | M | 53 | WBS | M | 46 |
| Control | M | 45 |  |  |  |
| Control | M | 19 | WBS | F | Teen |
| Control | F | 12 | WBS | F | 9 |
| Control | unknown | Teen |  |  |  |

**Table S2.**
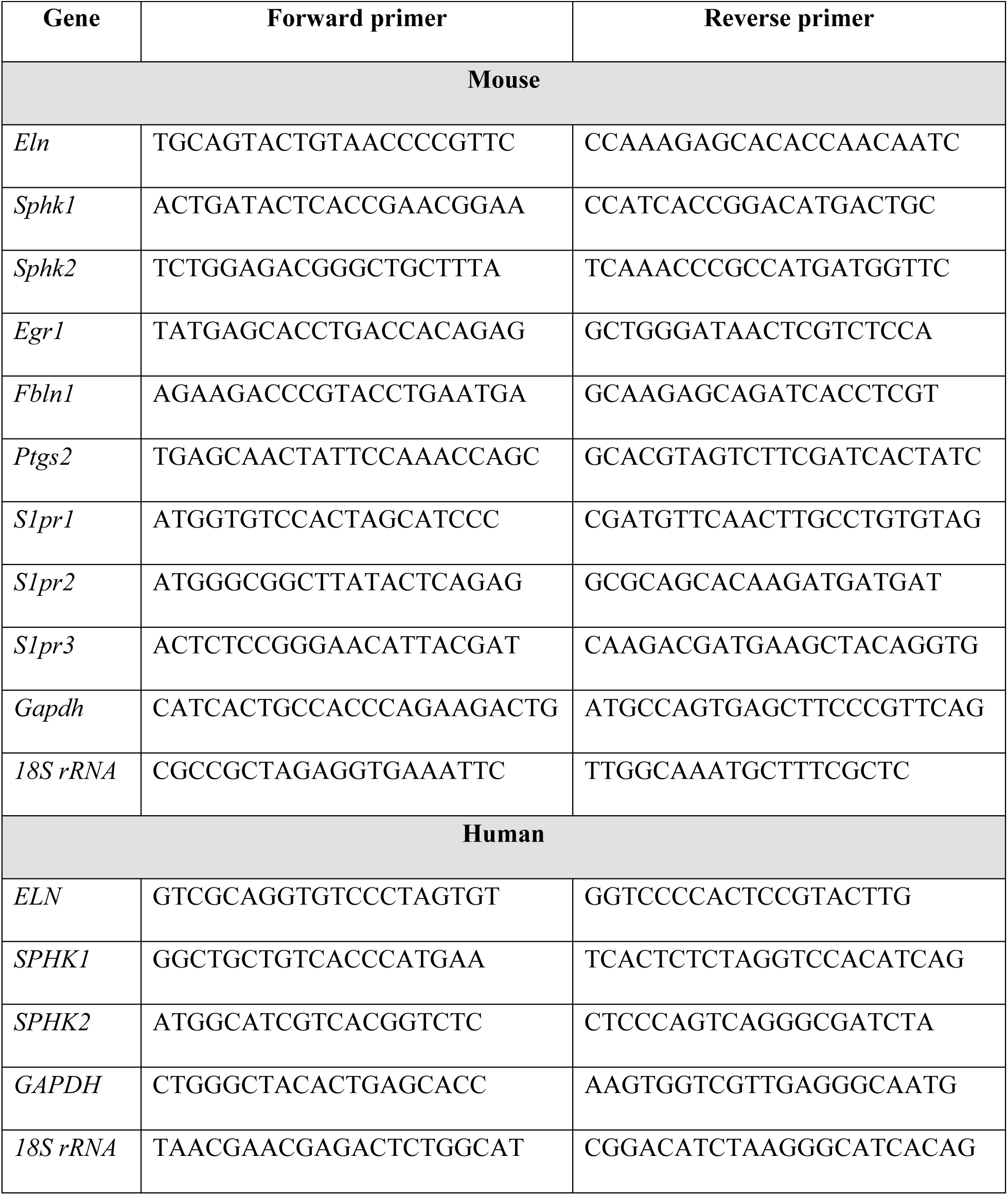
Primer pair sequences used for quantitative reverse transcription polymerase chain reactions.

**Table S3.**
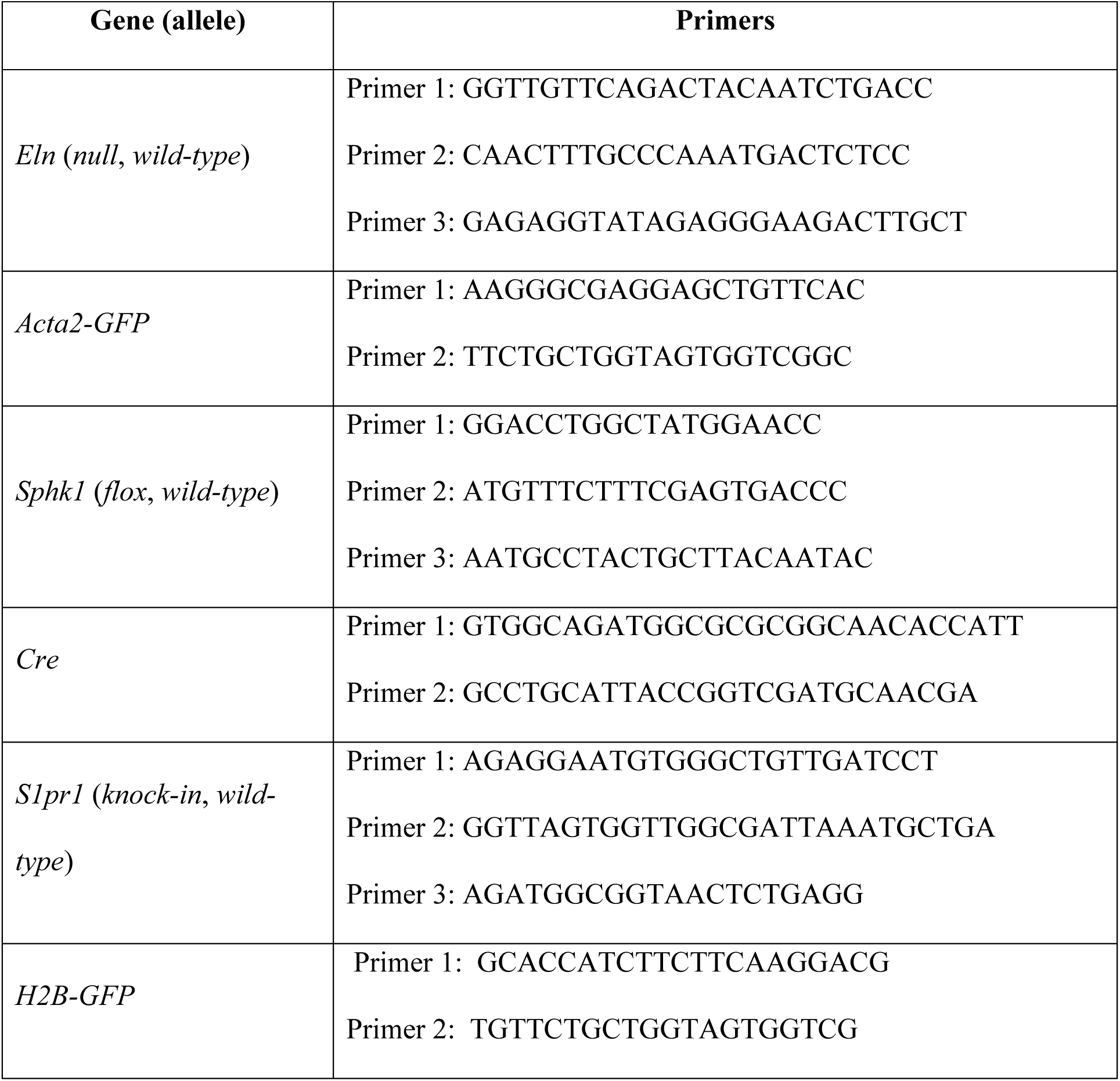
Primer sequences used for genotyping.

